# Critical role of Spatio-Temporally Regulated Maternal RNAs in Zebrafish Embryogenesis

**DOI:** 10.1101/2024.11.07.622483

**Authors:** Gopal Kushawah, Danielson Baia Amaral, Huzaifa Hassan, Madelaine Gogol, Stephanie H. Nowotarski, Ariel A. Bazzini

## Abstract

The maternal-to-zygotic transition shifts regulatory control from maternal to zygotic messenger RNAs (mRNA) through maternal mRNA degradation. While temporal aspects of maternal mRNA decay are known, spatial mechanisms remain underexplored. Using CRISPR-Cas9 and CRISPR-Cas13d systems, we functionally dissected the contribution of maternal versus zygotic fractions and overcame challenges of studying embryonic lethal genes. We identified differentially distributed maternal mRNAs in specific cells and evidenced the critical role of five maternal mRNAs, *cth1, arl4d, abi1b, foxa* and *lhx1a,* in embryogenesis. Further, we focused on the functionally uncharacterized *cth1* gene, revealing its essential role in gametogenesis and embryogenesis. *Cth1* acts as a spatio-temporal RNA decay factor regulating mRNA stability and accumulation of its targets in a spatio-temporal manner through 3’UTR recognition during early development. Furthermore, *Cth1* 3’UTR drives its spatio-temporal RNA localization. Our findings provide new insights into spatio-temporal RNA decay mechanisms and highlight dual CRISPR-Cas strategies in studying embryonic development.

**Highlights:**

1. Differentially distributed marginal maternal RNAs have a critical role in early embryogenesis.
2. Cas13d complements the Cas9 limitation to study the functions of embryonic lethal genes.
3. *Cth1* is essential for gametogenesis and early embryonic development.
4. *Cth1* is a maternal RNA decay factor required for spatio-temporal RNA regulation.
5. 3’UTR of *Cth1* drives its spatio-temporal RNA dynamics.

## Introduction

The maternal-to-zygotic transition (MZT) is a crucial phase in eukaryotic embryonic development, marked by the degradation of maternal mRNAs to enable zygotic genome activation (Vastenhouw et al., 2019; Walser and Lipshitz, 2011). In early development, maternal RNAs deposited by females before fertilization in oocytes controls the initial regulation. Their timely degradation is essential for normal embryogenesis, preventing developmental abnormalities. In zebrafish, mechanisms like microRNAs (e.g. miR-430) (Baia Amaral et al., 2024; Bazzini et al., 2012; Giraldez et al., 2006), codon optimality (Bazzini et al., 2016; Mishima and Tomari, 2016), M6A modifications (Zhao et al., 2017), and RNA-binding proteins have been identified as key regulators of maternal RNA decay (Despic et al., 2017; Mishima and Tomari, 2016; Zhao et al., 2020), ensuring the proper transition to zygotic control.

However, previously, most studies on maternal mRNA degradation have treated the whole zebrafish embryo as a homogeneous entity and used bulk mRNA sequencing techniques to study temporal changes (Baia Amaral et al., 2024; Bazzini et al., 2016; Giraldez et al., 2006; Vejnar et al., 2019; Zhao et al., 2017). While these approaches have advanced our understanding on temporal gene regulation, they could not reveal the functional importance of spatio-temporal maternal RNAs - RNAs that remain stable in specific cells of the embryo while being selectively degraded in other cells during early embryonic development. Recent single-cell RNA sequencing studies in zebrafish have shown the presence of maternal RNA signatures in early precursors of specific cell lineages during early development (Farrell et al., 2018; Wagner et al., 2018). Additionally, a recent study utilizing single-cell RNA SLAM-seq (Fishman et al., 2024) shows differential distribution of certain maternal RNAs in different cells during early development but did not account for the functional significance of such genes. Also, previous studies in zebrafish, *drosophila* and other animals have shown spatio-temporal RNA decay regulation of *Nanos* related gene and *Hsp83* gene (Gavis and Lehmann, 1992; Koprunner et al., 2001; Oulhen et al., 2013). Nonetheless, the functional implications of spatio-temporal maternal RNA localization resulting from differential cell-specific RNA degradation have not been thoroughly explored, primarily due to the absence of technologies capable of efficiently targeting maternal RNA during early vertebrate embryogenesis. Based on these observations, we hypothesize that spatio-temporal maternal RNA stability is essential for the proper differentiation and coordinated development of distinct cell types within the developing embryo.

Here, we first identified maternal RNAs with spatio-temporal patterns during early embryonic development. Then we focused on spatio-temporal maternal RNAs with marginal distribution and validated their patterns by RNA labelling during early embryonic development. Later, we used CRISPR-Cas13d RNA knockdown technology which is an efficient system for specific RNA decay during early embryogenesis (Bi et al., 2021; Buchman et al., 2020; da Silva Pescador et al., 2024; Hernandez-Huertas et al., 2024; Kim and Hutchins, 2024; Kushawah et al., 2020b; Liu et al., 2024; Morelli et al., 2023; Moreno-Sanchez et al., 2024; Treichel and Bazzini, 2022; Ventura Fernandes et al., 2024; Yang et al., 2024). CRISPR-Cas13d mediated knockdown of these genes displayed developmental phenotypes. The successful rescue of the knockdown phenotype validated the specificity of this system. Embryos injected with the CRISPR-Cas9 system did not recapitulate the early embryonic knockdown phenotypes, validating the minimal or no zygotic contribution in these phenotypes. From selected marginal spatio-temporal RNAs, we focused on the *cth1* gene. *Cth1* is among the highest maternally deposited RNAs with rapid spatio-temporal decay dynamics during embryogenesis and functionally uncharacterized in zebrafish (Fishman et al., 2024; Sur et al., 2023; te Kronnie et al., 1999). Different studies suggest that *cth1* is present in yeast and fishes and has its vertebrate as well as mammalian orthologs the Zinc Finger Protein 36 like (ZFP36l) genes (Makita et al., 2021; Martínez-Pastor et al., 2013; Stevens et al., 1998; Treguer et al., 2013). *Cth1* in yeast, along with its orthologs in higher vertebrates, has been shown to function as an mRNA decay factor by binding to AU-rich motifs in the 3’UTRs, playing an important role in various RNA metabolic processes (Barlit et al., 2024; Cicchetto et al., 2023; Martinez-Pastor et al., 2013; Vejnar et al., 2019). However, no functional studies have been conducted on *cth1* in zebrafish, apart from its potential AU-rich 3’UTR sequence binding capacity (te Kronnie et al., 1999; Vejnar et al., 2019). We found that *cth1* Cas9 genetic mutants fail to fertilize due to disrupted gametogenesis, leading to embryonic lethality. Therefore, CRISPR-Cas13d knockdown system complemented this limitation and elucidated the regulatory functions of *cth1* during early embryonic development such as affecting the mRNA stability and modulating the accumulation of mRNA targets by interacting with their 3’UTRs. Finally, we found that 3’UTR of *cth1* contains *cis* elements regulating its marginal spatio-temporal dynamics. Collectively, this study highlights the critical role of *cth1* as a key maternal spatio-temporal RNA decay factor essential for both embryogenesis and gametogenesis. Notably, this study is also the first to use both CRISPR-Cas13d and CRISPR-Cas9 systems to distinguish the functions of maternal versus zygotic contributions function in embryonic development. Further it demonstrates the ability of CRISPR-Cas13d in uncovering the functions of genes that are embryonic lethal or that leads to infertility.

## Results

### CRISPR-Cas13d knockdown of spatio-temporal maternal RNAs reveals their impact in embryonic development

We hypothesize that maternal RNAs can be degraded in a cell-specific manner during early embryogenesis, leading to a spatio-temporal distribution that may be essential for proper embryo development. To identify spatially distributed maternal mRNAs with low zygotic contribution, we combined single-cell spatial datasets (Satija et al., 2015) with SLAM-seq profiles (Baia Amaral et al., 2024). The SLAM-seq method differentiates between maternal and zygotic mRNAs by incorporating a UTP analog into newly synthesized mRNA transcripts (the zygotic component). This analog is detected as T -to- C mutations after the extracted RNA undergoes *in vitro* treatment prior to sequencing (Herzog et al., 2017b). Among the 8,920 genes with spatial and SLAM-seq data, 3,758 were maternally provided (CPMs > 20 at specific hours) with a low zygotic component (zygotic component < 0.1) (Figure 1A) (Baia Amaral et al., 2024; Bhat et al., 2023; Herzog et al., 2017a). By analyzing the median expression of marginal (meso-endodermal) and non-marginal regions per gene (Satija et al., 2015), a distinct set of genes with marginally localizing mRNAs was identified. This marginal region contains genes with a zygotic component (e.g. *gata5*) and a few that are mainly maternally provided (Figure 1B and Table S1). Within this group of marginal mRNAs, 5 genes displayed predominantly maternal deposition (Figure 1A - B). These five genes: *cth1, arl4d, abi1b, foxa* and *lhx1a* exhibited high unlabeled Counts Per Millions (CPMs) but low labeled CPMs during the maternal-to-zygotic transition (MZT), indicating that they are primarily maternally deposited mRNAs by 6 hours post fertilization (hpf) (Boxplot with line plot) (Figure 1C). To validate the spatial localization of these primarily maternal mRNAs in the marginal region of the embryo, Hybridization Chain Reaction (HCR) was performed at 6 hpf for *cth1, arl4d, abi1b, foxa* and *lhx1a*. As predicted, all candidate mRNAs were visualized in the marginal region, consistent with the spatial predictions (Figure 1D). In conclusion, these five genes are mainly maternally provided with low zygotic contribution and localized in the marginal region, suggesting that their localization may be due to differential stabilization or trafficking to the marginal region of the embryo rather than zygotic expression.

**Figure 1.**
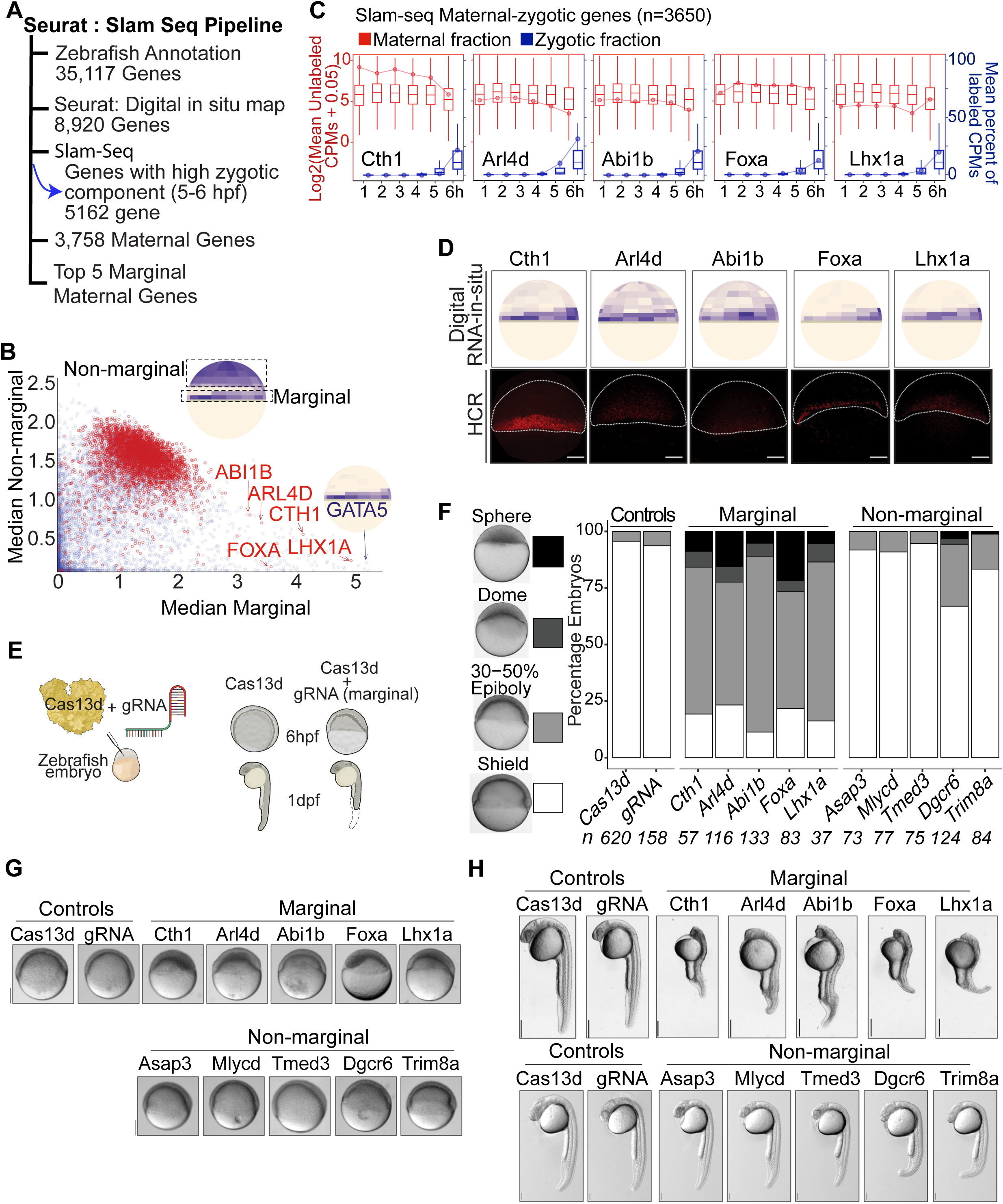
Identification, validation and functional analysis of spatio-temporal marginal maternal RNAs. A. Spatio-temporal maternal RNA selection pipeline: Data from Seurat: digital RNA- *in situ* prediction maps and SLAM-seq were combined to select marginal maternal RNAs. B. Scatter plot between median of marginal bin values and median of non-marginal bin values showing the differential distribution of different transcripts in embryo. Inset cartoon illustrated the bins selected for the calculation. Red and blue colored dots represent the pure maternal RNA transcripts and those with zygotic expression, respectively. The five selected marginal maternal RNAs *cth1, arl4d, abi1b, foxa* and *lhx1a* are indicated with red color while one marginal gene *gata5* with zygotic expression is shown by blue color. C. Plot showing the selected 5 marginal maternal RNA SLAM-seq time course and the predominance of maternal RNA fractions, as counts per million (CPMs) (red color) up to 6 hpf as compared to very low fraction of zygotic transcripts (blue color). D. Hybridization Chain Reaction (HCR) recapitulates the predicted digital RNA *in situ* patterns (up) of marginal maternal RNAs (*cth1, arl4d, abi1b, foxa and lhx1a*) at 6 hpf (down). Scale Bar, 100 μm. E. Schematic illustration of CRISPR-Cas13d knockdown assay: Cas13d protein along with cocktail of 3 gRNAs were injected into one cell zebrafish embryo. RNA knockdown phenotypes were observed at different developmental stages. F. Stacked bar plots showing the percentage of developmentally affected embryos for different targeted spatio-temporal maternal RNAs (marginal and non-marginal), along with controls (Cas13d and gRNA alone) at 6 hpf. The intensity of gray color is directly associated with severity of the developmental phenotypes (Sphere, dome, 30-50% epiboly, shield) respectively. G. Representative spatio-temporal mRNA knockdown phenotypes images of corresponding spatio-temporal maternal RNAs at 6 hpf along with controls (Cas13d and gRNAs alone, scale bar, 200 μm). H. Representative spatio-temporal mRNA knockdown phenotypes images at 1 dpf, Top panel targeted marginal maternal RNAs embryos and bottom targeted non-marginal maternal RNAs embryos along with corresponding controls (Cas13d and gRNAs alone). Scale bar top and down (1mm and 0.5 mm).

To find the functional significance of these spatio-temporal maternal RNAs during development, we knockdown these five marginal maternal RNAs as well as 5 non-marginal maternal RNAs as controls (Figure S1A) using the CRISPR-Cas13d knockdown system (Kushawah et al., 2020a). Three guide RNAs (gRNAs) against each candidate’s mRNA were designed and injected along with Cas13d protein into one cell stage zebrafish embryo separately for each gene (Figure 1E). Each targeted gene showed significant knockdowns assayed by qRT-PCR at 4 hpf as compared to controls (p-value ≤ 0.03, Cas13d alone vs Cas13d with each gRNA, t. test) (Figure S1C). To address the developmental effect of the knockdown during embryogenesis, the developmental stages were monitored at 4 hpf, 6 hpf and 26 hpf. While knockdown embryos had slight but significant developmental delay at 4 hpf as compared to the control (Cas13d) (p-value ≤ 1.2e-10, Cas13d alone vs Cas13d with each gRNA, chi-square test) (Figure S1B and S1D). At 6 hpf, the marginal maternal RNAs displayed strong developmental delay as compared to the controls (p < 2.2e−16, Cas13d alone vs Cas13d with each gRNA, chi-square test) (Figure 1E, 1F and S1E). In contrast, the knockdowns of most non-marginal gene did not display strong developmental delay at 6 hpf (p ≥ 0.34, Cas13d alone vs Cas13d with gRNA (*asap3, mlycd, tmed3*), p-value ≤ 1.8e-05, Cas13d alone vs Cas13d with gRNA (*dgcr6, trim8a*) chi-square test) (Figure 1F, 1G and S1E). At 26 hpf, broadly spatial context-specific phenotypes were observed (Figure 1H, S1F). Marginal gene knockdowns predominantly exhibit defects in body axis formation, particularly affecting notochord development (p-value ≤ 2.6e-05, Cas13d alone vs Cas13d with each gRNA, chi-square test) (Dal-Pra et al., 2011; Kamimoto et al., 2023), whereas only few non-marginal (anterior region) maternal RNA knockdowns (*dgcr6* and *trim8a*) primarily lead to abnormalities in brain development and slight developmental delay (p ≥ 0.07, Cas13d alone vs Cas13d with gRNA (*asap3, mlycd, tmed3, dgcr6*), p-value = 0.04, Cas13d alone vs Cas13d with gRNA (*trim8a*) chi-square test) (Figure 1H and S1F). Thus, all these results suggest that the marginally localized maternal mRNAs play critical roles during early embryonic development.

### CRISPR-Cas13d knockdown phenotypes are specific and can be rescued

To validate the specificity of knockdown phenotypes, rescue experiments were conducted. Specifically, new gRNAs were designed to target the 3’UTR of the endogenous genes. The rationale was to rescue the phenotype by co-injection of ectopic mRNA encoding the targeted gene but with non-targeting beta-globin 3’UTRs (Figure 2A). To conduct the rescue experiment, we first defined the overexpression mRNA concentration that would not compromise the development. For each candidate gene, 5ng/μl to 100ng/μl mRNA concentrations were injected in one cell stage embryos (5 to 100pg/embryo) (Figure 2B (25ng/μl), S2 A and S2B (5ng/μl to 100ng/μl)). For *abi1b, foxa* and *lhx1a*, we were able to determine the maximum injected mRNA concentration that did not phenotypically affect early development. However, for *cth1* and *arl4d,* embryos injected with even 5ng/μl displayed developmental abnormalities (Figure S2C).

**Figure 2.**
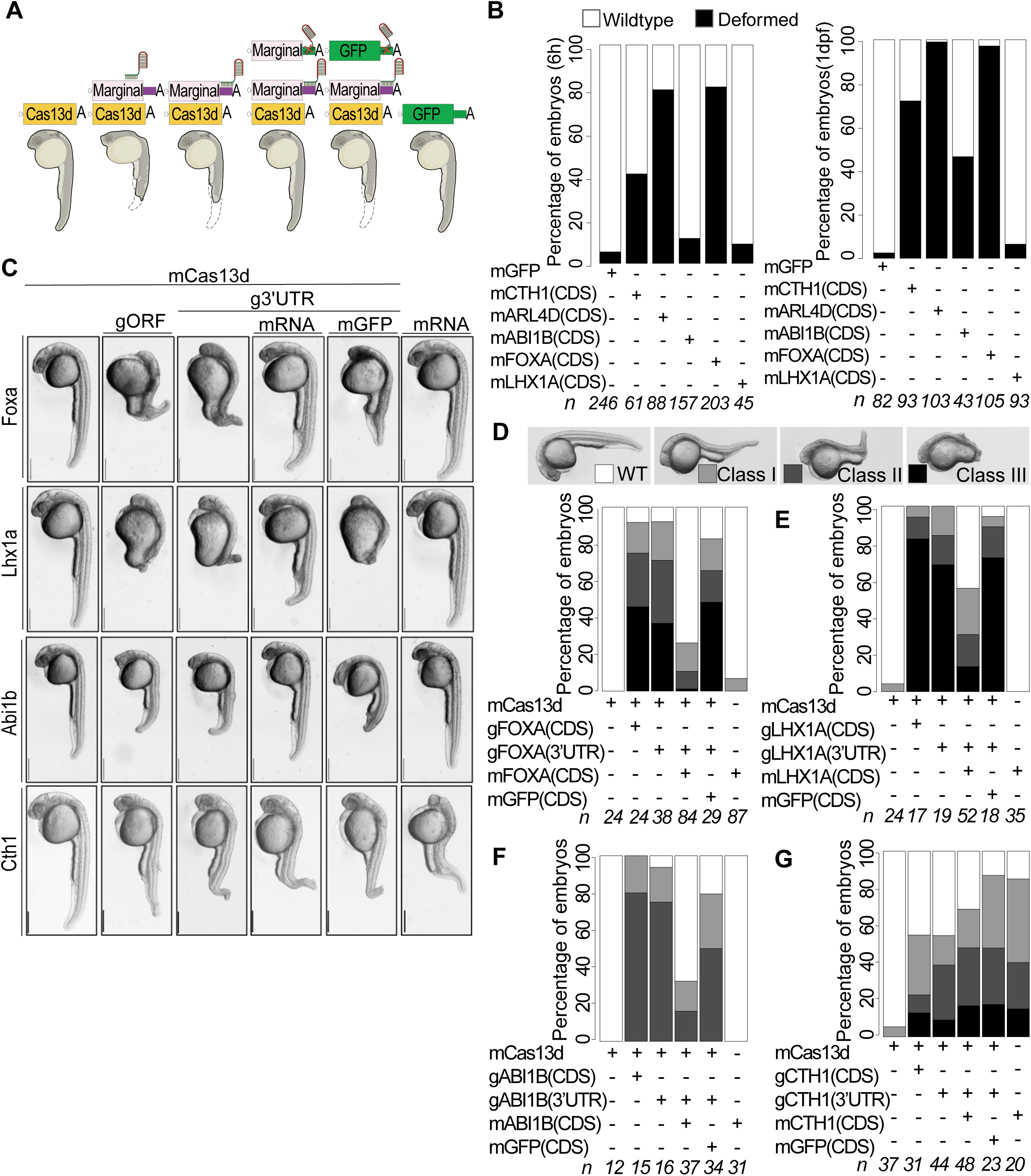
CRISPR-RfxCas13d mediated knockdowns are specific and can be rescued. A. Schematic illustration of Cas13d mediated RNA knockdown phenotype rescue assay in zebrafish. For rescue experiment, 3’UTR of endogenous transcript was targeted and ectopic RNAs from respective gene containing non-target 3’UTR was injected. Other panels show different kind of controls: Cas13d alone, Cas13d + gRNA targeting the ORF (gORF), Cas13d + gRNA targeting the endogenous 3’UTR + mGFP and mRNA alone of respective marginal gene). B. Stack bar plot showing the percentage of developmentally affected embryos after injecting with 25ng/μl of ectopic overexpression concentration for marginal genes (*cth1, arl4d, abi1b, foxa, lhx1a*) and 100pg/embryo for GFP as control at 6 hpf and 26 hpf. Black color represents developmental phenotype, and white color is for wildtype stage at given time points. n represents the number of embryos observed. C. Representative images for respective gene knockdown (Cas13d + g3’UTR), rescue panel (Cas13d + g3’UTR + mRNA) along with different knockdown (Cas13d alone, Cas13d + gORF) and rescue controls (Cas13d + g3’UTR + mGFP, mRNA alone). *Cth1* knockdown phenotype cannot be rescued, as its ectopic overexpression alone also show phenotypes. Concentrations of different mRNAs used in phenotype rescue Cas13d mRNA (300pg/embryo), gRNAs (500-600pg/embryo), rescue mRNA (*foxa* (5pg/embryo), *lhx1a* (75pg/embryo), *abi1b* (25pg/embryo), *cth1* (5pg/embryo), GFP mRNA (50pg/embryo) and ectopic mRNA alone concentrations are same as rescue mRNA for respective genes. Images are of 28 hpf embryos, scale bar, 1mm. D. Stack bar plots showing the percentage of observed phenotype rescue for *foxa* knockdown (D), *lhx1a* (E), *abi1b* (F) and *cth1*(G) at 1 dpf. Different intensities of gray color are positively associated with the severity of the phenotype (dark, most severe phenotype (class III) and white, (wildtype)). N represents the number of embryos observed in each condition.

Then, after defining the “no phenotype” overexpression concentration, the rescue experiments were performed. As expected, the co-injection of the gRNA targeting the 3’UTR of the marginal genes with mRNA encoding for Cas13d recapitulates the phenotype observed targeting the coding region of the same genes at 26 hpf, validating the specificity of the phenotype (Figure 2C). Remarkably, the co-injection of the overexpression constructs along with 3’UTR targeted knockdown injection mix significantly rescues the phenotype as early as 6 hpf for *foxa, abi1b* and *lhx1a* (*foxa* p-value < 2.7e-05, *lhx1a* p-value < 8.7e-11, *abi1b* p-value < 0.002, Cas13d alone vs rescue, chi-square test) as well as at 26 hpf for *foxa, abi1b* and *lhx1a* (*foxa* p-value < 1.4e-12, *lhx1a* p-value < 2.6e-05, *abi1b* p-value < 0.0001 (Cas13d alone vs rescue), chi-square test) (Figure 2C-G, S3A-E). The *cth1* phenotype was not rescued, however, with even the minimum injected overexpression concentration affecting embryonic development (Figure 2C, 2G, S2A-C, S3A, and S3E). And as expected, the ectopic expression of *gfp* did not rescue any knockdown phenotypes (*gfp* p-value ≥ 0.3, Cas13d alone vs gfp, chi-square test). Furthermore, the rescue experiment for *arl4d* was not performed because the minimal injected overexpression RNA concentration was either embryonic lethal or induced developmental deformities (Figure 2B, S2A-C). In sum, all these results together demonstrate that the rescuable CRISPR-Cas13d knockdown phenotypes are specific.

### Marginal maternal mRNA phenotypes are largely caused by the maternally deposited transcripts

To rule out that a minimum zygotic expression of selected marginal maternal candidate genes (Figure 1B) could explain knock down phenotypes at 6 hpf and 24 hpf, we targeted their zygotic DNA copies using the CRISPR-Cas9 system (Figure 3A) (Moreno-Mateos et al., 2015). Specifically, mRNA encoding for Cas9 was co-injected with a single or combination of three gRNAs targeting the coding region of each of the selected marginal candidate genes (*cth1, arl4d, abi1b, foxa* and *lhx1a*). First, as a proof of principle, we recapitulated the *slc45a2* gene (albino phenotype), specifically embryos co-injected with Cas9 and gRNA targeting *slc45a2* showed lack of pigmentation at 30 hpf compared to embryos injected with Cas9 alone (Figure S4A). Second, the mutagenesis rate of the marginal targeted genes was calculated at 6 hpf and 28 hpf by genotyping the targeted candidates by sequencing. All the loci targeted displayed more than an average of 84.5% and 86.3% mutagenesis rate at 6 hpf and 28 hpf, respectively (Figure 3B, 3C). At 6 phf, none of the embryos injected with Cas9 and with any gRNAs showed a developmental delay phenotypes compared to the control embryos (Cas9 alone, albino and non-injected) (p ≥ 0.30, Cas9 alone vs Cas9 plus each gRNA, chi-square test) (Figure S4B and S4C). At 30 hpf, no phenotype was observed after targeting the zygotic component for *cth1* (Figure 3D). For the other candidates*, arl4d, abi1b*, *foxa* and *lhx1a*, few of the embryos displayed a very mild body axis deformity as compared to the Cas13d knockdowns phenotypes when their maternal and zygotic components were targeted (p = 0.02 to 1.4e-13, (Cas9 alone vs each marginal, chi-square test) (Figure 3D, 3E and S4D). These results indicate that the five (*cth1, arl4d, abi1b, foxa and lhx1a*) marginal maternal mRNAs are strongly contributing to these developmental phenotypes and with no major role of potential zygotic components during early embryogenesis.

**Figure 3.**
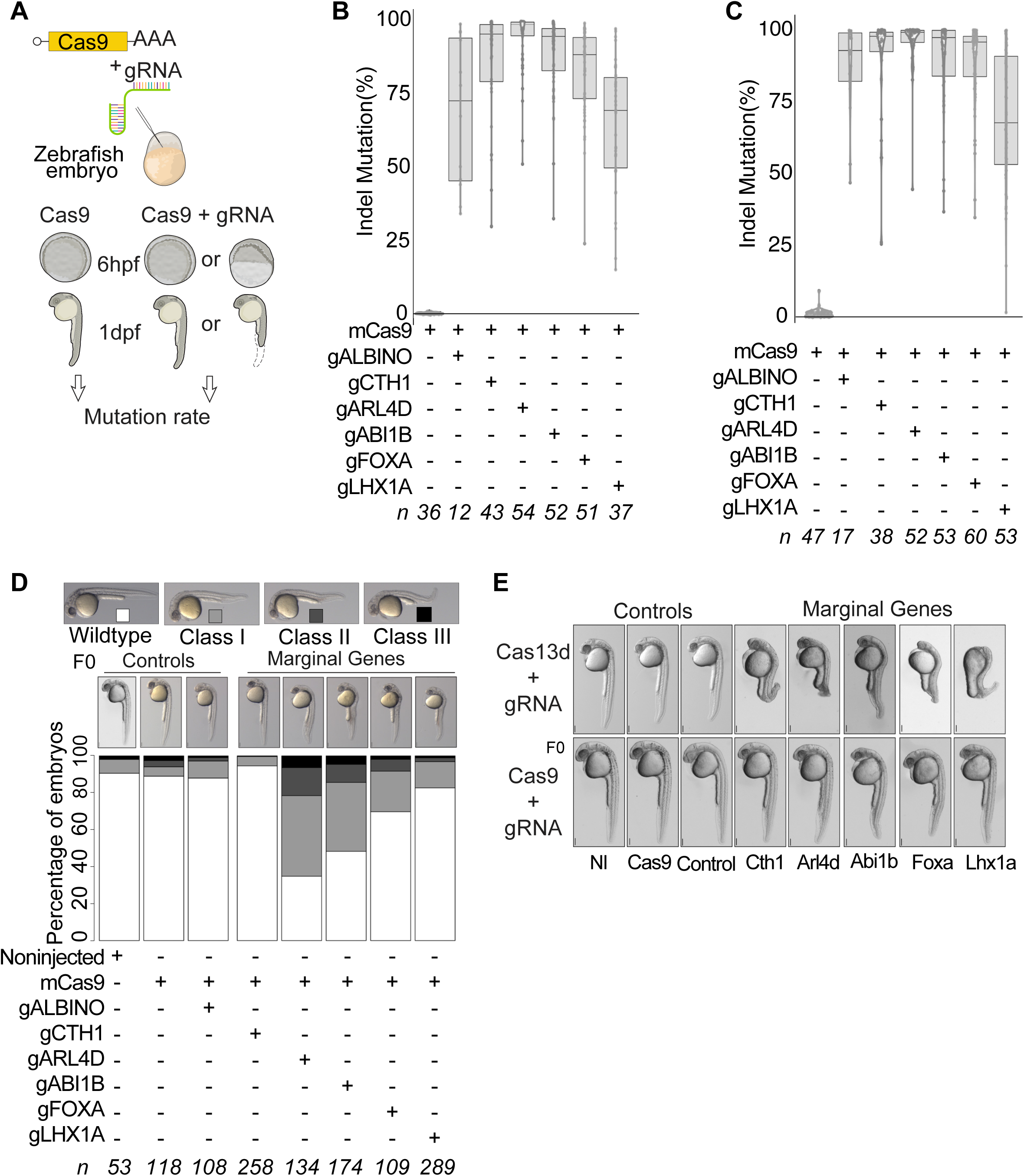
CRISPR-Cas9 mutation of marginal genomic DNA show minimal or no zygotic contribution to early developmental phenotypes. A. Schematic illustration of CRISPR-Cas9 mutation assay in zebrafish: 100ng/μl of Cas9 mRNA and 20ng/μl of each gRNA was injected at one cell embryo and phenotypes were followed at 6 hpf and 1 dpf. B. B. and C. Density box plot showing percentage of high levels of Cas9 indel mutation frequencies for associated zygotic copies of marginal genes at 6 hpf (B) and 1 dpf (C) embryos. n represents the numbers of embryos sequenced for each gene at 6 hpf and 1 dpf. The average indel mutation frequency for marginal genes at 6 hpf (84.5%) and 1 dpf (86.3%). D. Representative images and stack bar plot for percentage of Cas9 mediated indel mutation phenotypes of zygotic copies of marginal genes (*cth1, arl4d, abi1b, foxa, lhx1a*) at 1 dpf along with different controls (noninjected, Cas9 alone, *albino*). Different intensities of gray color are directly associated with severity of developmental phenotype (Class III (most severe) and white (wildtype). n represents the number of embryos observed. Scale bar, 0.5mm. E. Representative images of 1 dpf embryos showing comparison of Cas13d mediated marginal gene transcripts knockdown phenotypes (Top) vs Cas9 mediated indel mutagenesis in zygotic copies of marginal genes. Scale bar, 0.5mm.

### Cth1 mutants are infertile and have abnormal gametes

From all selected marginal maternal candidates, *cth1* emerged as an interesting candidate as its functions are unexplored in zebrafish development. *Cth1*, in yeast and its mammalian orthologs is an RNA-decay factor binding to ARE-rich sequences in the 3’UTR and has a critical role in various vital developmental processes (Cicchetto et al., 2023; Lai et al., 2003). In zebrafish, *cth1* is one of the most highly deposited but rapidly decaying mRNAs during early embryogenesis (Figure 4A) (Fishman et al., 2024). *Cth1* transcripts show spatio-temporal decay dynamics during early embryonic development (Figure 4B) (Sur et al., 2023). Both overexpression and knockdown of *cth1* affect development, making the knockdown phenotype not rescuable (Figure 1, 2). Moreover, *cth1* F0 independent mutants, generated with Cas9 showed a high mutagenesis rate but do not exhibit early developmental phenotypes (Figure 3) suggesting that the early phenotype was caused by the maternal contribution (Figure 1 and 3). Interestingly, *cth1* F0 mutants generated were apparently phenotypically normal adults (Figure 4C), however exhibited infertility problems (Figure 4D, S5A). These infertility phenotypes were observed in the mutants with each of the three gRNA individually and when combined, indicating these problems are unlikely due to off-target effects (Figure 4D, S5A). Specifically, unviable or degraded embryos resulted from crossing F0 mutants (both male and female) with wild-type or within *cth1* mutants (Figure 4D). Furthermore, both oocytes and sperm from all independent lines were defective, displaying from 80 - 100% completely degraded and unviable oocytes (Fig 4E, S5B). Oocytes from these mutant females were translucent white, with ruptured membrane as compared to opaque pale yellow, intact and viable wildtype mature oocytes (Figure 4E). Mature sperm squeezed from mutant males were also more translucent white as compared to opaque milky white liquid in wildtype sperm (Figure S5C). In addition, mature sperm from mutant males have abnormal phenotypes including rough and degraded sperm heads as compared to smooth, membranous wildtype sperm heads (Figure 4F). Moreover, mutant sperm show tail abnormalities ranging such as no tail, short tail, two tails, serrated tail and multiple tails with a completely ruptured membrane sheath (Figure 4F and S5D) and are unable to fertilize wildtype embryos (Figure 4D and S5A, S5E). These results highlight the crucial role of *cth1* in fertility, oogenesis and spermatogenesis. Mutations in *cth1* orthologs in other animal models have been reported to cause female infertility (Ball et al., 2014; Ramos et al., 2004; Zhou et al., 2023) but here we show that in zebrafish, *cth1* also has a functional role in male spermatogenesis which was not reported for its orthologs in other animal models, suggesting its critical role in producing healthy gametes and maintaining the reproductive health of both male and female zebrafish.

**Figure 4.**
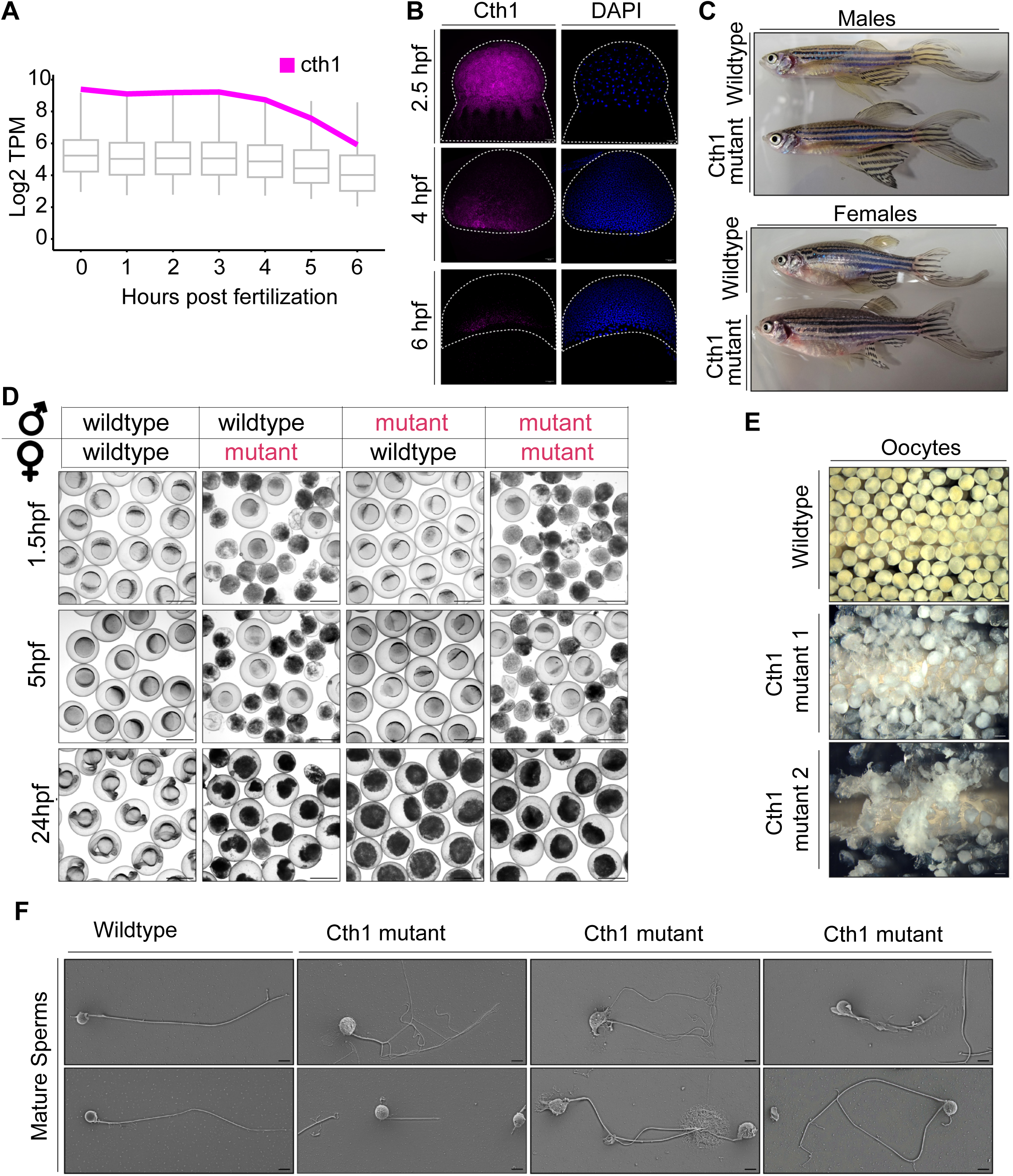
*Cth1* Cas9 mutants are infertile with abnormal mature gametes. A. Boxplot showing total mRNA level (RNA-seq) time course line graph from 0 to 6 hpf in wildtype conditions. *Cth1* transcript expression is shown, it is strongly maternally deposited and show rapid decay kinetics. B. Hybridization Chain Reaction (HCR) experiment showing *cth1* transcript spatio-temporal dynamics from 2.5 hpf (uniform distribution) to 6 hpf (marginal distribution). *Cth1* transcript is labeled with magenta color and blue color is for DAPI nuclear stain. C. Representative phenotype images of adult *cth1* mutant fish (Cas9) with their corresponding wildtype controls. Top panel is for males, bottom panel is for females. All the fish are 14 months old. D. Representative images of fertilized embryos from Adult Cas9 *cth1* F0 mutants (red color text) bred with same age wildtype fish (black text), or mutant counterpart (both red text) produce either dead (mutant female) or unfertilized embryos (mutant male), while control embryos are viable. (Panels show 1.5 hpf, 5 hpf and 24 hpf embryos). Scale bar, 1000 μm. E. Representative images of mature oocytes from two independent *cth1* Cas9 mutant females as compared to oocytes from wildtype female of same age (14 months old) are translucently white, fragmented and degraded. Scale bar 1000 μm. F. Representative scanning electron images of mature sperm from wild type and *cth1* mutant males: Mature sperm from wildtype males (14 months old) have normal morphology with one rounded smooth head and one tail. While *cth1* mutant sperm from independent mutant males (14 months old) have rough, wrinkled sometimes degraded head and tail with different morphologies ranging from no tail, short, to multiple fragmented tails phenotypes. Scale bar is 3 μm.

### Cas13d uncovers Cth1 regulatory functions during early embryonic development by complementing the CRISPR-Cas9 system limitations

As Cas9 *cth1* mutants were infertile, we capitalized on the CRISPR-Cas13d technology to unravel the regulatory function of *cth1* during early development. First, we confirmed that no extra band is present between 28s and 18s rRNA in the *cth1* knockdown background, confirming no CRISPR-Cas13d induced collateral activity (Figure S6C) (Kushawah et al., 2020a; Moreno-Sanchez et al., 2024). Then RNA-seq was performed in CRISPR-Cas13d mediated *cth1* knockdown compared to embryos injected with Cas13d alone at 6 hpf (Figure 5A). *Cth1* showed a 1.56-fold reduction (adjusted p value = 1.13e-09) in expression and Gene Ontology (GO) analysis at 6 hpf suggests that *cth1* is associated with genes involved mainly in DNA maintenance, energy metabolism, angiogenesis, hematopoietic process, and vesicle mediated transport to membrane (Figure S6A). *Cth1* and its orthologous gene family are suggested to bind to AU rich sequences in the 3’UTR of transcripts (Cicchetto et al., 2023; Hu et al., 2020; Martinez-Pastor et al., 2013; Vejnar et al., 2019), therefore, we selected common candidates with ARE motifs in the 3’UTR that were upregulated in the *cth1* knockdown and/or were reported as targets of *cth1* orthologs in different animals by previous studies (Barlit et al., 2024; Cicchetto et al., 2023) (Figure 5A). Further, single-cell RNA seq data (Sur et al., 2023) suggests that the spatio-temporal expression patterns of these potential targets are mutually exclusive with respect to the expression pattern of *cth1* during early embryonic development (Figure 5B).

**Figure 5.**
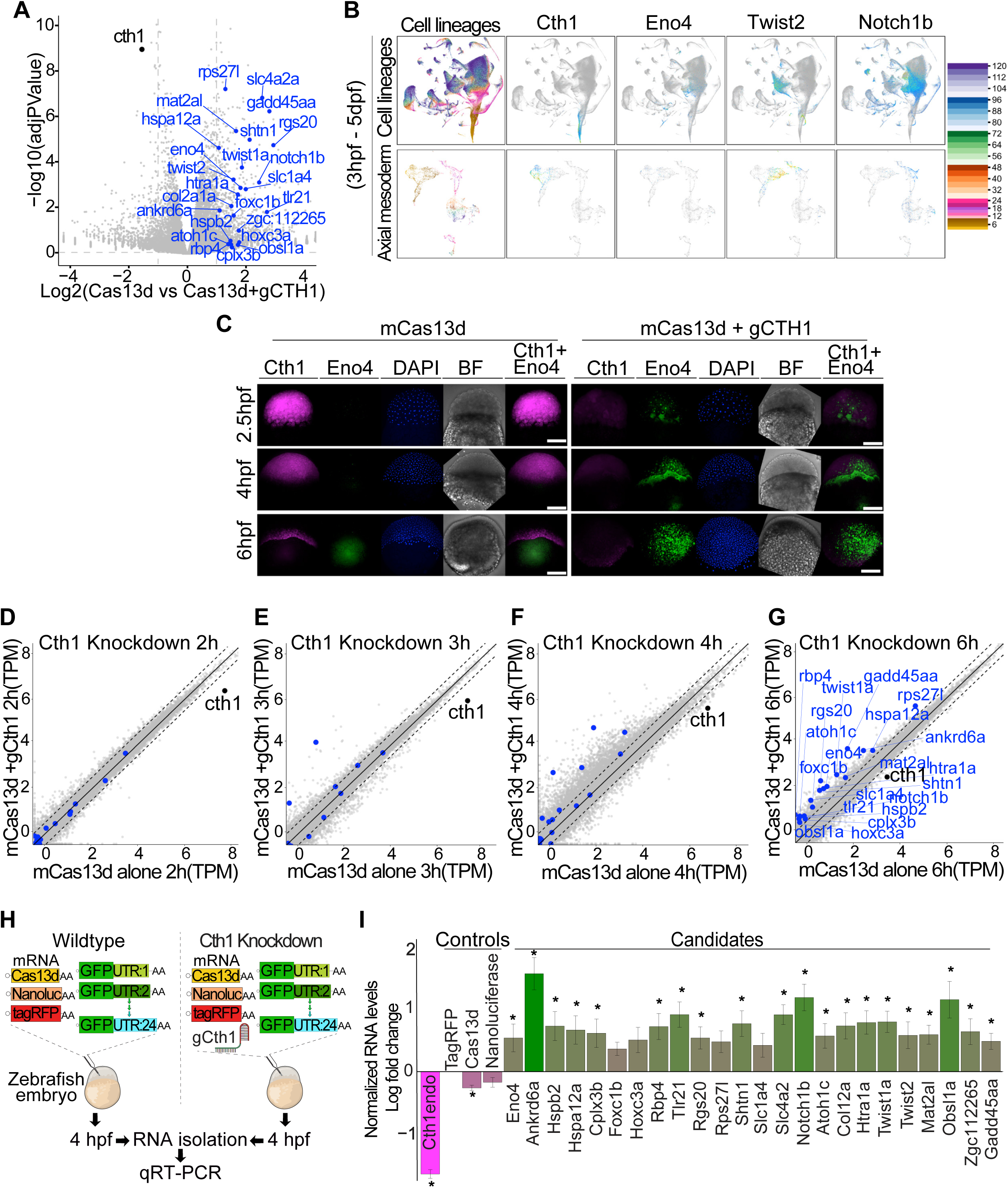
CRISPR-Cas13d uncover the functions of embryonic lethal Cth1 gene during development. A. Volcano plot showing *cth1* transcript (black dot) knockdown as compared to embryos injected with Cas13d alone (log fold change 1.56-fold reduction and adjusted p value = 1.13e-09). Potential *cth1* targets from its ortholog studies in different animal models and cell lines as well as other upregulated candidates with “AU” rich sequences in their 3’UTRs were highlighted in blue. B. Single cell RNA seq trajectory data using *Daniocell,* an online zebrafish single cell RNA-seq resource from 3hpf to 5dpf with different cell lineages and axial mesodermal cell lineages. *Cth1* trajectory plot showing the presence of *cth1* transcripts only at early developmental stages (up to 6-7 hpf) and its expression in axial mesoderm. Trajectory data from its potential targets is (*eno4, twist2, notch1b*) showing the accumulation or presence of their transcripts once *cth1* transcripts disappear, in a mutually exclusive manner in different cell lineage from 3hpf to 5dpf. Below panels show distribution in axial mesodermal cell lineage (3hpf to 5dpf). C. HCR in wildtype and Cas13d mediated *cth1* knockdown conditions for *cth1*(magenta), *eno4* (green) along with DAPI and bright field images at 2.5, 4, and 6 hpf. Upon *cth1* knockdown, *eno4* transcripts start accumulating at early developmental stages. Scale bar, 100 μm. D. Schematic illustration of GFP:3’UTR reporter assay in wildtype and Cas13d mediated *cth1* knockdown conditions followed by qRT-PCR. 3’UTR from 24 different potential targets (10pg/embryo each) were cloned downstream of GFP, and different injection controls like mRNAs from tagRFP (10pg/embryo), nano luciferase (10pg/embryo), and cas13d (300pg/embryo) were included in this assay. Embryos with this small library was injected in one cell zebrafish embryo with and without gRNA (600pg/embryo) against *cth1.* RNA at 4hpf from both conditions were isolated for qRT-PCR. E. The qRT-PCR plot showing normalized levels of RNA log fold change between embryos injected with the cocktail of mRNA with vs without the gRNA targeting *cth1* at 4 hpf. Endogenous levels of *cth1* transcripts were decreased (magenta color) in the embryos containing both cas13d and the gRNA compared to the embryos without gRNA. As expected, the injections controls (Cas13d mRNA, nano-luciferase levels) displayed similar relative level compared to tagRFP in the embryos injected with or without the gRNA targeting *cth1*. But the GFP:3’UTRs of all the potential targets were upregulated in the *cth1* knocked-down embryos (green color). Significant p value < 0.05, from at least 5 biological replicates from two independent experiments (t. test). F. Different scatter plots (F, G, H and I) showing Cas13d mediated *cth1* knockdown RNA seq time course at 2 (F), 3 (G), 4 (H) and 6 hpf (I) highlighting *cth1* and its potential targets. mRNA levels in *cth1* knockdown conditions (y-axis) vs control, Cas13d alone (x-axis) were plotted at 2, 3, 4 and 6 hpf. *Cth1* mRNA is indicated by black color and its potential targets by blue color. From 2 to 6 hpf the levels of *cth1* potential targets increase, suggesting their upregulation upon *cth1* knockdown. At least 2 biological replicates were used in all the conditions at different time points.

To investigate the effect of *cth1* knockdown on the early onset of its spatio-temporal dynamics*, in situ* HCRs for *cth1* and its potential target *eno4* in zebrafish embryos were conducted at various developmental stages in the *cth1* knockdown and in the control embryos. Interestingly, *eno4*, shows early transcript accumulation in *cth1* knockdown embryos, compared to the control embryos (Figure 5C). These findings suggest that *cth1* regulates the spatio-temporal dynamics of its potential target transcripts during early embryonic development.

Therefore, to examine the kinetics of *cth1* targets regulation, RNA-seq was performed on knockdown embryos (Cas13d and single *cth1* guideRNA) and control embryos (Cas 13d alone) at 2-, 3-, 4-, and 6-hours post-injection. Interestingly, most of the potential gene targets*’* transcript levels showed an early and consistently higher accumulation in the *cth1* knockdown embryos (Figure 5F - I). These findings are consistent with our *in-situ* results (Figure 5C) and suggest that *cth1* may delay the accumulation of these target transcripts during early embryogenesis.

Consequently, we hypothesized that the knockdown of *cth1* may cause an early onset of transcription of target mRNA or delay their degradation. To investigate the latter option, a mixture of 24 mRNA encoding for GFP with respective 3’UTRs of the potential *cth1* targets was co-injected along with tagRFP, Cas13d and nano-luciferase mRNAs as internal controls, with or without the gRNA targeting *cth1*, into one-cell stage zebrafish embryos (Figure 5D). Interestingly, the GFP:3’UTR mRNAs were relatively more stable than the internal control mRNA (tagRFP, nano luciferase, and Cas13d mRNA) in the *cth1* knockdown embryos as compared to the control injected embryos (Figure 5E, S6B), suggesting that *cth1* regulates mRNA stability by recognizing the 3’UTRs of its targets. Moreover, iCLIP data indicated that *cth1* preferentially binds to 3’UTR sequences (Vejnar et al., 2019). And altogether, these results suggest that *cth1* may prevent the accumulation of these targets, implying that while these targets are transcriptionally activated after zygotic genome activation, their accumulation is prevented by *cth1*-triggered decay (Figure 5C, F, G, H, I).

### *Cth1* RNA contains *cis*-regulatory elements in its 3’UTR driving the marginal localization

Several lines of evidence indicate that the localization of *cth1* mRNA in the marginal region of the embryo at 6 hpf is not driven by cell-specific transcription. *Cth1* is primarily maternally provided, with minimal zygotic transcription detected via SLAM-seq (Figure 1). *Cth1* is one of the most highly maternally deposited mRNAs but is also very unstable during MZT (Figure 4A, 4B). Therefore, we hypothesized that the spatio-temporal localization of *cth1* mRNA is likely regulated by cell-specific mRNA degradation or protection mechanisms during embryogenesis. To investigate this, we co-injected *in vitro* transcribed mRNAs encoding GFP with the 3’UTR of *cth1*, along with TagRFP as an internal control, into single-cell zebrafish embryos. The localization of GFP and TagRFP was identified using HCR at 2, 4, 5 and 6 hours post injection (Figure 6). Interestingly, GFP mRNA containing the *cth1* 3’UTR localized to the marginal region of the embryos at 4 hpf, whereas TagRFP control mRNA was uniformly distributed across the embryos (Figure 6). These findings strongly indicate that the *cth1* 3’UTR harbors cis-regulatory elements that dictate its localization to the marginal region during early embryogenesis.

**Figure 6.**
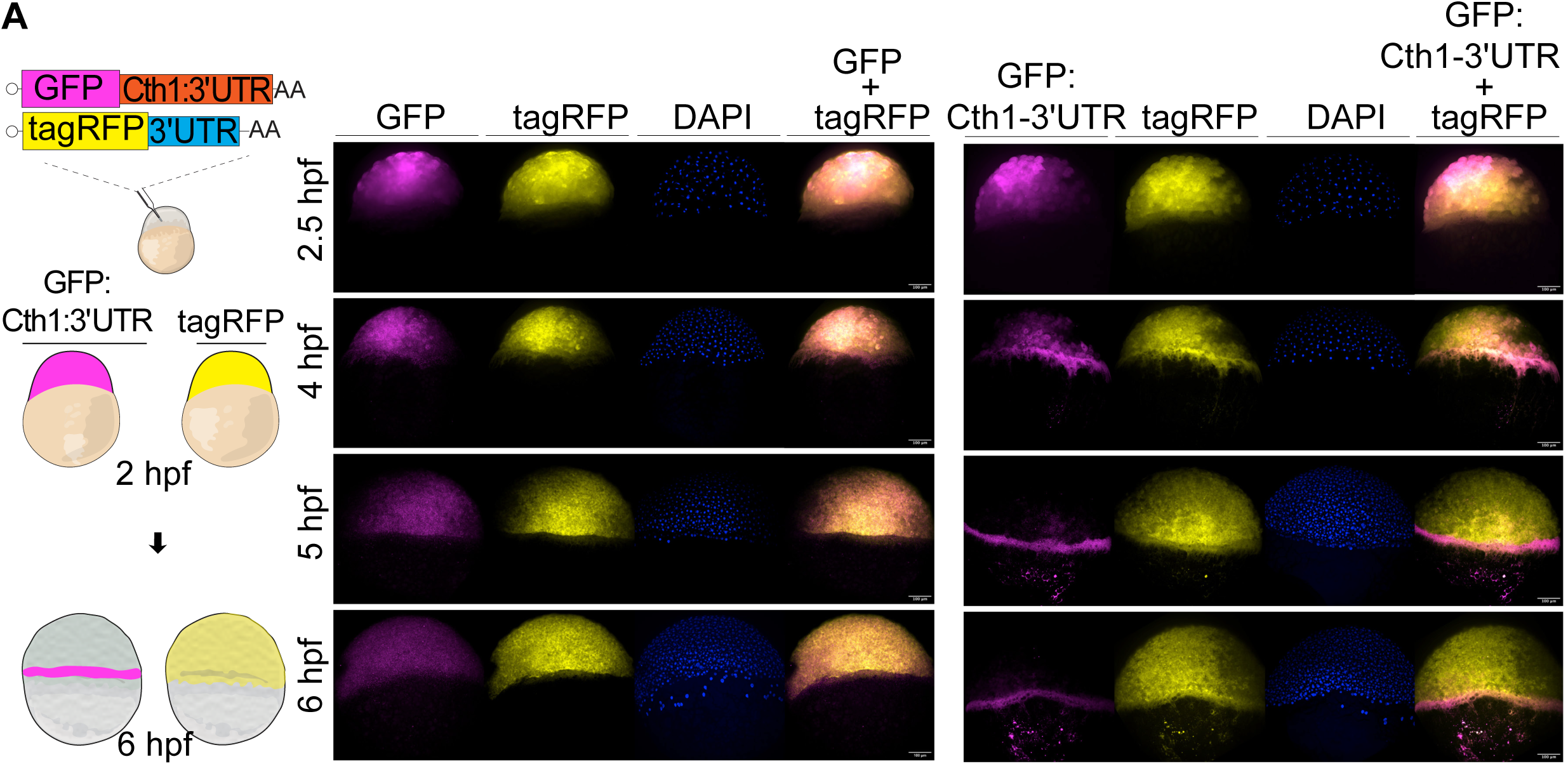
*Cth1* 3’UTR is regulating spatio-temporal localization. A. Cartoon showing injection strategy for HCR against injected GFP: *Cth1* 3’UTR along with control tagRFP at 2 and 6 hpf. HCR for control GFP and control TagRFP in first panel, while for GFP: *Cth1* 3’UTR along with control tagRFP in second panel at 2.5, 4, 5 and 6 hpf. Magenta color is for GFP RNA, yellow color is for tagRFP RNA, Blue color is for DAPI stain, and composite images are for GFP and TagRFP. Scale bar, 100 μm.

## Discussion

We have uncovered that at least five maternal mRNAs are spatio-temporally regulated (Figure 1) and their knockdowns (Figure 1) and in some cases their ectopic expression (Figure 2) affects early embryonic development. Previously, spatio-temporal regulation was shown mainly for germ cell related genes, such as *nanos* (Koprunner et al., 2001), however this type of regulation now appears to be more widespread. And while, in this study we focused on *cth1*, the successful knockdown phenotype rescue experiments of *foxa, abi1b* and *lhx1a* (Figure 2) provide an innovative system to further study their functions during early embryogenesis in the future. Furthermore, there are more maternal transcripts localized in the marginal region that can also be investigated (Figure 1B). And while this work is focused on differentially distributed maternal RNAs in the marginal region, spatio-temporal maternal RNA regulation from other regions like anterior, posterior, dorsal, ventral, and many other specific part of the embryo will be interesting to explore in future (Satija et al., 2015).

In this study, we have introduced a pioneering use of CRISPR-Cas9 and CRISPR-Cas13d systems to unravel the distinct contributions of maternal and zygotic components during early embryonic development. By utilizing CRISPR-Cas13d to simultaneously knock down both maternal and zygotic transcripts, independently followed by targeted disruption of zygotic DNA using CRISPR-Cas9 (Figure 1 and Figure 3), we provide a precise framework for delineating the role of maternal and zygotic fractions in developmental processes. These parallelly independent yet complementary approaches not only enhance our ability to accurately dissect these complex contributions but also significantly improve the efficiency of experimental procedures, optimizing both time and resources. The CRISPR-Cas13d approach becomes essential when genetic mutants are embryonic lethal (Figure 7). Here, Cas13d effectively complemented this limitation, as demonstrated in our study that *cth1* Cas9 mutant led to infertility problems, and CRISPR-Cas13d revealed the essential functions of *cth1* during early embryonic development. In the future, this dual CRISPR-Cas system approach will be very useful in uncovering the functions of embryonic lethal genes in different animal models.

**Figure 7.**
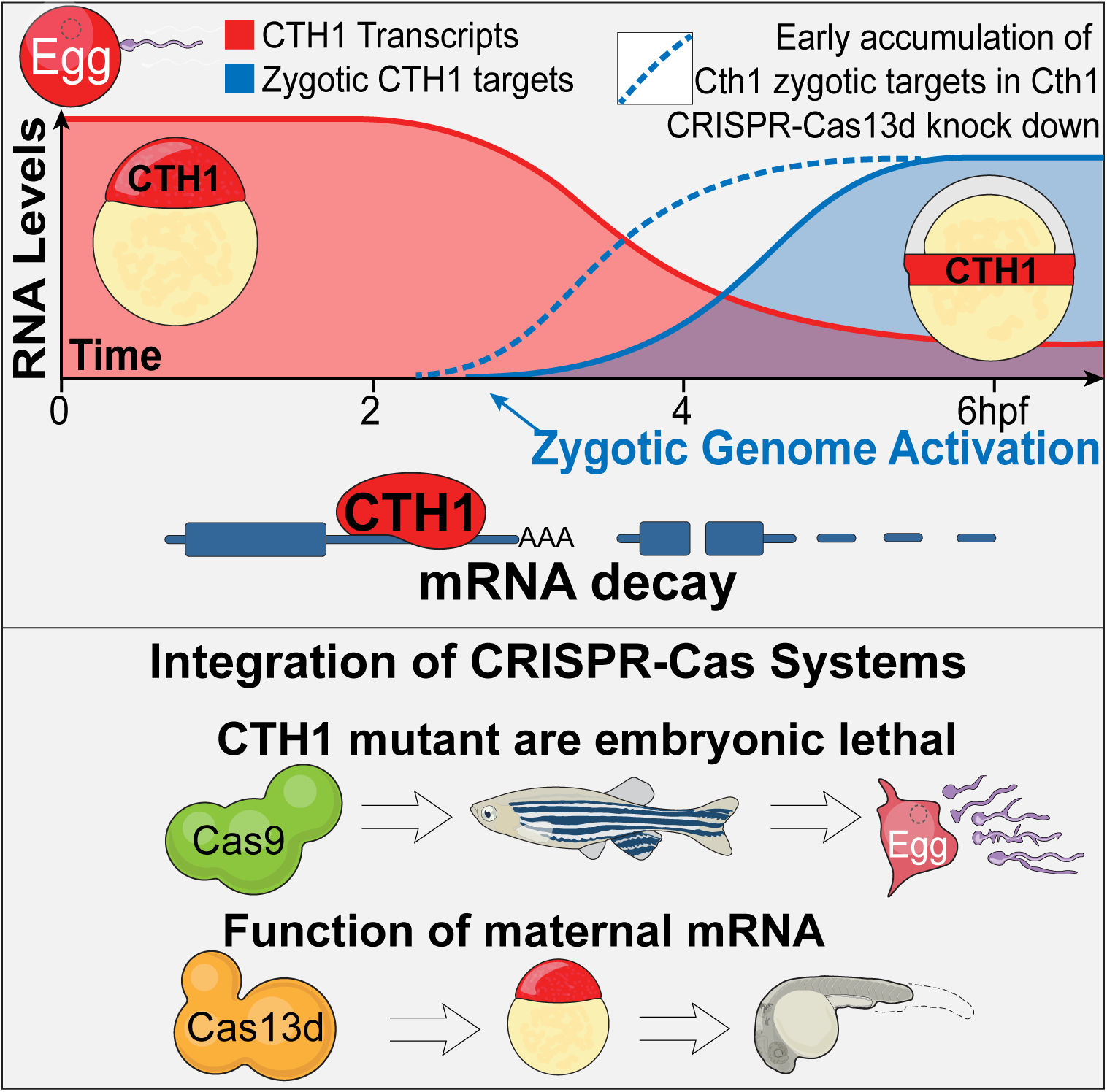
*Cth1* is a highly expressed, maternally provided mRNA that undergoes degradation during the maternal-to-zygotic transition and accumulates in the marginal region of zebrafish embryos. Knockdown of *cth1* using the CRISPR-Cas13d system disrupts embryonic development and delays the accumulation of potential mRNA targets by affecting their stability in a 3’ UTR-dependent manner. CRISPR-Cas9 *cth1* mutants were infertile, highlighting the essential role of *cth1* in oogenesis and spermatogenesis. CRISPR-Cas13d allowed us to study the *cth1* function during embryogenesis, bypassing the embryonic lethality observed in mutants.

Interestingly, all the *cth1* ortholog genes reported in other animal models, show female-specific fertility problems (Ball et al., 2014; Khalaj et al., 2016; Ramos, 2012; Treguer et al., 2013; Zhou et al., 2023) but in our study, *cth1* is also shown to disrupt male gametogenesis (Figure 4F). To advance our understanding, future research will focus on elucidating the specific role of *cth1* in gametogenesis and understanding how these mutations impact the overall health of adult mutants. Mutations in human ortholog of *cth1* gene and other vertebrate orthologs were linked to embryo implantation failure (Ball et al., 2014; Ramos, 2012; Ramos et al., 2004; Zhou et al., 2023), therefore this study could also provide a strong ground for human fertility research. Although the adult mutant fish apparently did not exhibit observable developmental phenotype abnormalities, in the future it will be interesting to investigate the internal morphology of tissues, especially given the severe gametogenesis defects observed (Figure 4). Careful investigation of the testis, ovary, and other closely related organs in these mutants could reveal interesting *cth1* biology and its implication in human health.

From the molecular point of view, our reporter assays in *cth1* knockdown conditions have shown that *cth1* is a spatio-temporal RNA decay factor regulating the stability of transcripts containing “AU” rich motifs in the 3’UTR, which is further supported by iCLIP data (Vejnar et al., 2019) and its orthologs*’* role in other animal models (Cicchetto et al., 2023; Cook et al., 2022; Lai et al., 2003; Ramos et al., 2004) (Figure 5). This reporter assay also provides an innovative approach to use the CRSPR-Cas13d system to unravel mRNA stability independent of *de novo* transcription of the potential targets and control analyzed (Figure 5). Interestingly, the fact that the *cth1* targets analyzed are mostly zygotically expressed (Figure 5D, 5F-5G); and in the *cth1* knockdown, they show increased stability (Figure 5A, 5C, 5D, 5F - 5I), as well as higher accumulation at early time points (Figure 5C, 5F - 5I), suggest that *cth1* targets might be transcriptionally active but fail to accumulate due to the *cth1* mediated RNA decay (Figure 7). Recently, it was shown that miR430 can also prevent or delay the accumulation of zygotic miR430 targets (Baia Amaral et al., 2024). Therefore conceptually, it can be proposed that the zygotic genome might get activated in a much broader sense from the transcription point of view, however, the post-transcriptional gene regulation mechanisms (e.g. miR430 and *cth1*) shape or prevent the temporal accumulation of several mRNAs. Single cell RNA-seq data also suggested that *cth1* potential targets do not share same spatio-temporal context during development as their transcripts are either distributed in different cells or show delayed onset of transcription in the same cell lineage during zebrafish development (Figure 5B) (Farrell et al., 2018; Sur et al., 2023). Thus, based on our results, *cth1* emerged as a spatio-temporal maternal RNA decay factor regulating the stability and therefore accumulation of zygotic mRNAs during the maternal to zygotic transition in zebrafish embryos (Figure 7). In the future, it will be interesting to dissect the molecular mechanism (e.g. deadenylation), targets site recognition, define all its targets and to interrogate its function in other species during embryogenesis.

Finally, it is intriguing how *cth1* mRNA gets localized to the marginal region. *Cth1* is one of the most abundant maternal mRNAs after fertilization, however it is very unstable (Figure 4a) with a low zygotic component (Figure 1). The combination of SLAM-seq and single cell RNA-seq data has suggested that the small zygotic component of *cth1* is preferentially coming from the marginal region (Fishman et al., 2024). However, most of the *cth1* mRNA at 6 hpf is maternal (Figure 1) and the genomic mutagenesis did not compromise early development (Figure 3). Also, more interestingly, the injection of an mRNA reporter containing the *cth1* 3’UTR was localized to the margin region, suggesting that the *cth1* 3’UTR drives the localization of the maternal mRNA (Figure 6). While we cannot rule out any intercellular trafficking of the mRNA, the instability of the *cth1* mRNA suggests that its cell-specific degradation might cause this marginal localization. Two main fundamental questions raised from our results that need to be addressed in the future are: Which are the *cis* regulatory elements in the *cth1* 3’UTR driving the marginal localization? And What are the factors and mechanism affecting the stability of *cth1* mRNA or the other factors in a cell-specific manner during embryogenesis?

## Limitations

This study has identified many interesting maternally provided differentially distributed RNA candidates, but in this study, we focused on *cth1* in detail. In the future, the authors would like to explore the functional regulation of the remaining maternally provided marginal as well as other differentially distributed maternal candidates during early embryogenesis. This is the first functional study for *cth1* functions in any vertebrate, and the authors would like to explore its loss of function mutants in detail to understand the detailed biology of *cth1* during gametogenesis and early embryogenesis. And further studies are needed to find out *cth1* binding motifs in 3’UTR sequences of its potential targets and compare it with its orthologs in other animals. Finally, we need to find out the sequences of *cis*-regulatory elements in the 3’UTR of *cth1*, responsible for its spatio-temporal dynamics during early embryogenesis.

## Lead Contact

Further information and resource request should be directed to lead contact Ariel A. Bazzini.

## Data and Code Availability

The Accession number for the RNA-Seq data used in this manuscript is GEO: GSE281036. All the relevant data is available upon request from the corresponding author.

## Experimental Model and Subject Details

### Zebrafish Maintenance, embryo productions and experimentations

All zebrafish (*Danio rerio*) experiments were performed in accordance with protocols approved by the Institutional Animal Care and Use Committee (IACUC) at the Stowers Institute. Adult zebrafish from the AB, TF, and TLF strains, aged between 6 to 18 months, were used for embryo production. Fish were randomly selected from a colony of approximately 500 individuals, consisting of four independent strains. For microinjections, at least 12 male and 12 female zebrafish were randomly bred to get fertilized timed one cell stage embryos. Embryos were maintained in zebrafish embryo media at 28.5°C under standard laboratory conditions.

## Method Details

### Hybridization chain reaction (HCR)

Candidate transcripts to label were selected, and corresponding probes, amplifiers, and reagents were ordered from Molecular Instruments. On the subsequent day, amplifiers tagged with specific fluorophores were added during the amplification step, followed by overnight incubation at room temperature in the dark. Finally, the embryos were washed and nuclear DAPI stain was added before imaging.

### Zebrafish microinjections

All the experiments were done by injecting one cell stage dechorionated zebrafish embryos. All the injected controls and experiment embryos were grown at 28.5°C using E2 zebrafish embryo media with .01% methylene blue. For Cas13d-mediated knockdown experiments, 300pg of Cas13d mRNA or 2.5-3ng of Cas13d protein was injected per embryo. Guide RNA (gRNA) concentrations for Cas13d knockdowns ranged from 400-600pg. For Cas9 mutagenesis, 100pg of Cas9 mRNA and 25pg of each gRNA were used per injection. The specific concentrations of transcripts and injection mixtures for various reporter assays and qRT-PCR experiments are detailed in their respective method sections.

### CRISPR-Cas13d mediated RNA knockdowns in zebrafish embryos

We employed the CRISPR-Cas13d system to achieve RNA knockdown in zebrafish embryos, as described previously (Hernandez-Huertas et al., 2022; Kushawah et al., 2020a). For the design of guide RNAs (gRNAs), we utilized the online RNA fold server (Hernandez-Huertas et al., 2022). To target specific spatio-temporal maternal RNAs, we designed a minimum of three gRNAs for each target RNA. Either Cas13d protein (3 ng/embryo) or Cas13d mRNA (300pg/embryo) and 3X gRNAs (500-600pg/embryo) (Table S2) for each RNA were injected at one cell stage zebrafish embryo. Following microinjections, all embryos were treated in similar manner, frequent cleaning, and incubated at 28.5 degrees Celsius. In general, at least 20 embryos for each replicate were collected for both qRT-PCR and RNA-seq. During phenotype quantification experiments, a cocktail of 3 gRNAs mix was used, but for knockdown rescue experiments and Cth1 RNA-Seq, reporter assay, only one gRNA targeting *cth1* was used. During this study, embryos were periodically cleaned and imaged at 4 hpf, 6 hpf and at 1-day post-fertilization (dpf).

### RNA isolation and qRT-PCR

To further test the knockdown efficiency and find the molecular signatures of knockdowns, RNA was isolated using either the standard TRIzol method (ThermoFisher Scientific) or the Zymo-Research RNA Mini Kit (ZYMO RESEARCH, Cat. No. R1055). The concentration of the isolated RNA was measured using either a NanoDrop spectrophotometer or the Qubit RNA-Broad Range assay. For qRT-PCR analysis, complementary DNA (cDNA) was synthesized according to the manufacturer’s protocol using the SuperScript IV First-Strand Synthesis System (ThermoFisher Scientific, Cat. No. 18091050) and using different oligos mentioned in table (Table S2). Quantitative real-time PCR (RT-PCR) was then performed using the low ROX SYBR-Green dye (Quanta Bio, Cat. No. 95074-05K), gene-specific primers (20μM), and 1-2μl of cDNA diluted in a 25μl reaction volume, depending on the gene of interest and the developmental time point. The reactions were conducted on an automated Freedom EVO® PCR workstation (Tecan) using a predefined template program designed for the number of samples. For normalization, endogenous housekeeping genes such as cdk2ap2 or taf15 were used as reference genes for target gene expression analysis, while for reporter assays, other injection controls like tagRFP, Nano Luciferase, or Cas13d were utilized.

### Knockdown phenotype rescue

To assess whether Cas13d-mediated knockdowns are specific and can be rescued, guide RNAs (gRNAs) were designed to target the 3’UTR of the gene of interest (Table S2), following a previously published protocol (Hernandez-Huertas et al., 2022). To recapitulate the knockdown phenotype, gRNAs specific to the open reading frame (ORF) of the target gene were used. For rescue experiments, we complemented the endogenous RNA by cloning its ORF together with a non-target beta-globin 3’UTR (Table S2), allowing for the expression of the transcript without interference from the gRNA targeting the endogenous 3’UTR. As a negative control, we utilized a GFP construct containing the beta-globin 3’UTR, which is not targeted by the gRNA against the respective gene 3’UTR, serving also as a control for any ectopic overexpression phenotype. For microinjection into one-cell zebrafish embryos, in vitro transcription of the gRNAs and other constructs was performed according to published protocols (Hernandez-Huertas et al., 2022; Kushawah et al., 2020a). In these experiments, gRNA concentrations ranged from 400 to 500pg per embryo, while 300pg of Cas13d mRNA was injected. The concentrations of the rescue transcripts varied depending on the specific marginal genes (*foxa* = 5pg, *abi1b* = 25pg, *lhx1a =* 70pg and *cth1 =* 5pg), with identical concentrations used for the corresponding negative rescue controls, GFP with non-target beta-globin 3’UTR. The same concentrations were also applied to ectopic expression controls in each respective set of rescue experiments.

### CRISPR-Cas9 mediated mutation in zebrafish

In this strategy, we designed 3 gRNAs (Table S2) targeting the zygotic genes of the corresponding spatio-temporal maternal RNAs. gRNAs were designed using CRISPR Scan guidelines (Moreno-Mateos et al., 2015). The Cas9 plasmid was digested using *XbaI* enzyme and capped *in vitro* transcription was performed using mMESSAGE mMACHINE™ T3 Transcription Kit to generate Cas9 mRNA (Vejnar et al., 2016). The gRNAs were synthesized according to the protocol outlined for the AmpliScribe-T7 Flash Kit. Zebrafish embryos at the one-cell stage were injected with either 100pg of Cas9 mRNA and 25pg of each of the three gRNAs in combination or with 25pg of each gRNA separately. After injection all the embryos were processed and incubated similarly. Phenotypes of the injected embryos were observed and imaged at 6 hpf and 1 dpf. During this process individual embryos were collected at 6 and 24 hpf in QuickExtract™ DNA Extraction Solution (Lucigen) for Indel mutations genotyping using MiSeq sequencing.

### Scanning Electron Microscopy

Sperm samples were diluted onto coverslips coated with 4nm of gold palladium with a Leica EM Ace600 and prepared as previously described for preserving membranes (Korneev et al., 2021). After critical point drying with a Tousimis Samdri-795 and coated with an additional 4nm of gold palladium sperm were imaged at 300kV, 100pA with an SE2 detector on a Zeiss Merlin SEM. All the images were processed similarly in Fiji using CLAHE plugin.

### Mature oocytes and sperm collection

To collect gametes from both wild-type and mutant zebrafish adults, we adhered to the institutional animal handling guidelines. For the collection of mature oocytes, female fish were housed with males in the same tank overnight, separated by a divider. The following morning, upon the activation of the light cycle, females were retrieved, anesthetized using 4 g/L MS-222, and stripped to collect the oocytes. For the collection of mature sperm, male fish were anesthetized using the same procedure. The anesthetized males were positioned in a fish holder with their ventral side facing upwards, and sperm was collected using a microcapillary pipette attached to an aspiration tube assembly.

### Image acquisition

Different microscopes were used according to the requirement. For zebrafish embryos grouped pictures at early time point up to 1 dpf Leica stereo-fluorescent microscope with DFC900 camera. For individual representative images Leica MZ APO stereo microscope was used for bright field images. For HCR images Nikon spinning disc a confocal microscope were used at 10x and 20x magnification for different fluorophores with same setting and exposure times for controls and respective experimental samples. For large adult fish Samsung galaxy s21 ultra camera was used. For sperm images Zeiss Merlin SEM with SE2 detector was used. For image processing Fiji software was used and image panels were made using adobe illustrator 2024.

### RNA-seq libraries preparation and Analysis

Cas13d mediated knockdown at 6 hpf: High quality RNA once passed through Bioanalyzer (Agilent) with RIN score more than 7.5 were used for library preparation. At least 500ng of RNA was used for library preparation using manufacturer’s protocol for the NEBNext Ultra II Directional RNA prep kit for Illumina (Cat. No. E7760S) with NEBNext Multiplex Oligos for Illumina (96 Unique Dual Index Primer Pairs (Cat. No. E6440). Further libraries were again checked on Bioanalyzer (Agilent) and Qubit (Life Technologies). Libraries were then pooled, quantified and sequenced as single read on a NextSeq 500 with high output read length of NextSeq-HO-75 on illumina flow cell (NextSeq 2000 P2 v3 (100 cycles), Cat. No. 20043738).

*Cth1* knockdown RNA-seq time course: library preparation was done using reagents from Watchmaker Genomics, for library preparation Watchmaker mRNA Library Prep Kit (Cat. No. 7BK0001) with xGen Stubby Adapter and UDI Primers (IDT, Cat. No. 1000592) were used. Samples were sequenced using paired reads on G4-F3 with a read length of 50bp using a flow cell from Singular Genomics (Cat. No. 700125). RNA-seq reads were demultiplexed into fastq format allowing up to one mismatch, using Illumina bclconvert (v 3.10.5) for samples sequenced on the NextSeq 500, and Singular Genomics sgdemux (v 1.2.0) for samples sequenced on the G4-F3. Subsequently, the reads were aligned to *GRCz11* reference genome from Ensembl using STAR (v 2.7.10b). The gene model retrieved from Ensembl, release 106 was used to generate gene read counts. The transcript abundance TPM (Transcript per Million) was quantified using RSEM (v 1.3.1).

Differentially expressed genes were determined using R package edgeR (v 3.42.4). Prior to differential expression analysis, low-expression genes were filtered out based on a cutoff of 0.2 CPM (Counts Per Million) in at least one library. The resulting p-values were adjusted with Benjamini-Hochberg method using R function p.adjust. Genes with an adjusted p-value < 0.05 and a fold change of 2 were considered as differentially expressed.

### Gene Ontology (GO) analysis

Gene functional enrichment analysis or GO analysis was performed using a custom script built over the R package clusterProfiler (v4.4.4). *GRCz11* Gene–GO terms retrieved from Ensembl BioMart were used to identify overrepresented GO terms in the differentially expressed genes compared to the background list of all expressed genes.

### Reporter assays in zebrafish embryo

To construct a GFP - 3’UTR library, the 3’UTRs of potential targets identified from RNA-seq data were amplified using cDNA isolated from 6 hpf zebrafish embryos. The amplified 3’UTRs were individually cloned downstream of GFP and confirmed by sequencing. Positive clones were digested with the NotI-HF enzyme, followed by *in vitro* transcription (IVT) using the SP6 mMESSAGE mMACHINE Maxiscript kit. Subsequently, an RNA cocktail was prepared, comprising 24 different GFP-3’UTR reporter transcripts, each at a concentration of 10 ng/μl. To assess the stability of these GFP-3’UTR reporters, the RNA cocktail, along with various injection controls including tagRFP (10ng/μl), nanoluciferase(10ng/μl), and Cas13d mRNA (300 ng/ μl), was injected into one-cell-stage zebrafish embryos under both wild-type and *cth1* knockdown conditions. For *cth1*knockdown, gRNA (600ng/μl) targeting *cth1* was co-injected alongside the injection mix used for wild-type conditions. Following injections, embryos were collected at the 4hpf stage for RNA extraction, subsequent cDNA synthesis, and qRT-PCR analysis.

### Quantification and Statistical analysis

To select different sample sizes, no predefined statistical method was used. The experiments were not randomized, and investigators were not blinded to group allocation during the experiments or outcome assessment. All collected data were included in the analysis without exclusion. All the quantification, gene expression and RNA-Seq plots were generated in R. Phenotype quantification across different CRISPR-Cas-mediated injection conditions was analyzed using chi-squared (χ²) tests, performed using R. For quantitative RT-PCR analysis, p-values were determined using an unpaired t-test, and error bars represent the Standard Error of the Mean (SEM). As mentioned in the RNA-seq methodology, fold changes for each gene were calculated using edgeR (version 3.42.4), following the exclusion of genes with a count of less than 0.2 CPM (Counts Per Million) in at least one library. P-values were subsequently adjusted using the Benjamini-Hochberg method, implemented in R, with statistical analysis performed on at least two biological replicates.

### Declaration of generative AI and AI-assisted technologies in the writing process

During the preparation of this work the authors used ChatGPT in order to improve the text. After using this tool, the authors reviewed and edited the content as needed and take full responsibility for the content of the publication.

## Acknowledgments

We thank Miguel A. Moreno-Mateos for critical reading and valuable suggestions for this manuscript. Carrie Carmichael for zebrafish related discussion and training. Different core facilities at Stowers Institute like Sequencing and Discovery Genomics, genome engineering, automation and PCR technology, Media prep and all members of the Bazzini laboratory for their intellectual and technical support. This study was supported by the Stowers Institute for Medical Research. AAB was awarded a Pew Innovation Fund and with the US National Institutes of Health (NIH-R01 GM136849 and NIH-R21 OD034161). This work was performed as part of post-doctoral training for GK at the Stowers Institute for Medical Research. This work was performed as part of thesis research for D.B.A. at the Graduate School of the Stowers Institute for Medical Research. Original data underlying this manuscript can be accessed from the Stowers Original Data Repository at http://www.stowers.org/research/publications/libpb-2515.

## Author contributions

G.K. and A.A.B. conceived the project and designed the research.

G. K. performed all zebrafish experiments

DBA performed the SLAM-Seq experiment and analyzed the data.

M.G. extracted Seurat digital *in situ* images.

H.H. did primary RNA-seq analysis.

G.K, DBA and A.A.B. performed data analysis.

S.N. performed electron microscopy.

and G.K, and A.A.B. wrote the manuscript with input from the other authors. All authors reviewed and approved the manuscript.

---

**Supplementary Figure 1.**
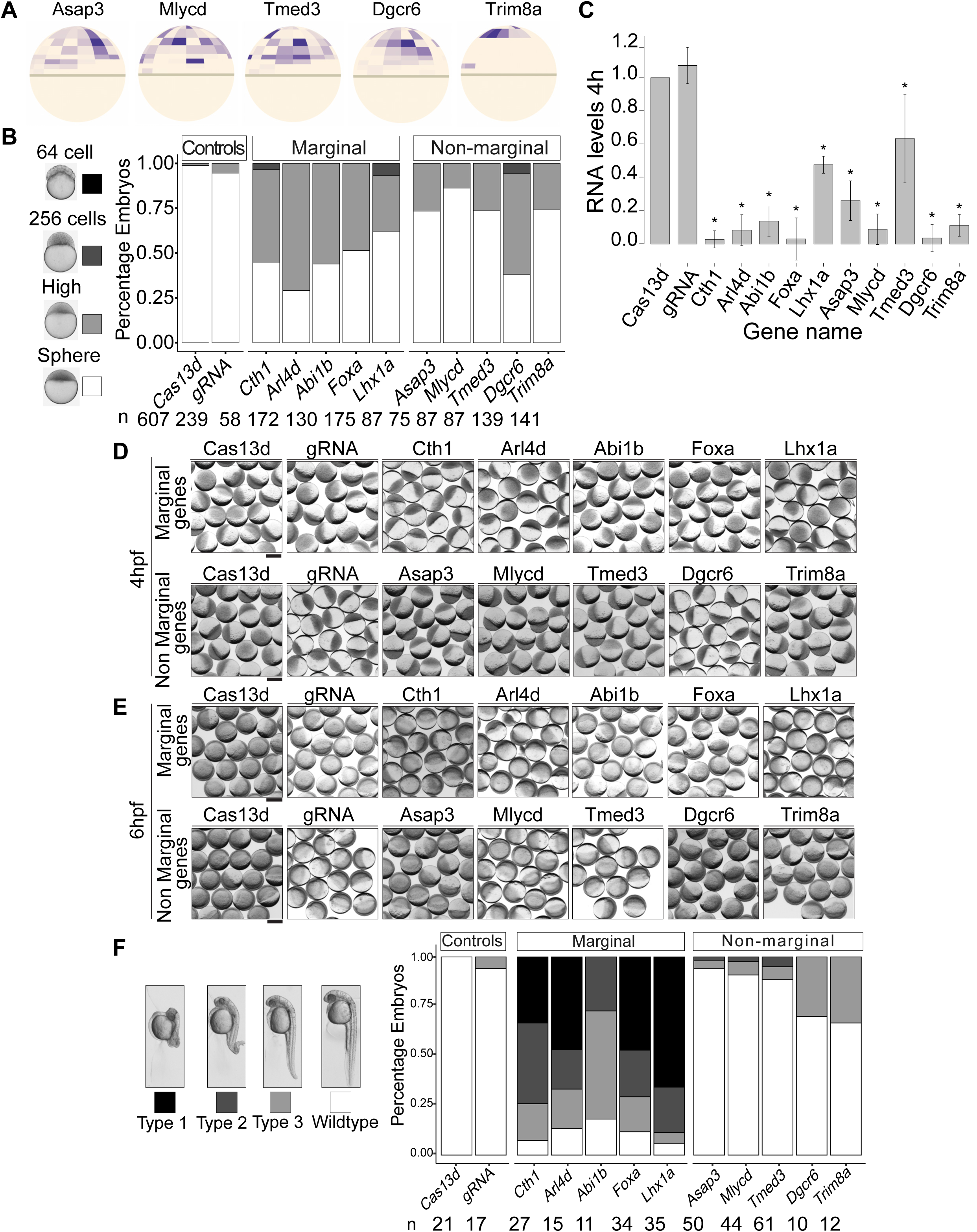
Critical role of spatio-temporal maternal RNAs in early embryogenesis. A. Digital RNA *in situ* for non-marginal controls (*asap3, mlycd, tmed3, dgcr6, trim8a*). B. Stacked bar plots showing the percentage of developmentally affected embryos for different targeted spatio-temporal maternal RNAs (Marginal and non-marginal), along with controls (Cas13d and gRNA alone) at 4hpf. Different intensities of gray color are directly associated with severity of developmental phenotype (64 cells, 256 cells, high, sphere) respectively. n represents the number of embryos observed at 4hpf. C. Bar-plot showing RNA levels upon CRISPR-Cas13d mediated knockdown (marginal and non-marginal RNA) at 4hpf measured by qRT-PCR. All the targeted RNA shows significant level of RNA knockdowns, (p-value ≤ 0.03, t. test). D. E. CRISPR-Cas13d knockdown representative phenotype images of corresponding spatio-temporal maternal RNAs at 4 (D) and 6 hpf (E) for marginal and non-marginal transcripts along with different controls (Cas13d alone and gRNA alone). Scale bar,100μm. F. RNA knockdown phenotype quantification at 26 hpf: Stacked bar plots with the percentage of developmentally affected embryos for different targeted spatio-temporal maternal RNAs (Marginal and non-marginal), along with controls (Cas13d and gRNA alone) at 26 hpf. Different intensities of gray color are directly associated with severity of developmental phenotype (Type 1, Type 2, Type 3, Wildtype) respectively. n represents the number of embryos observed at 26 hpf.

**Supplementary Figure 2.**
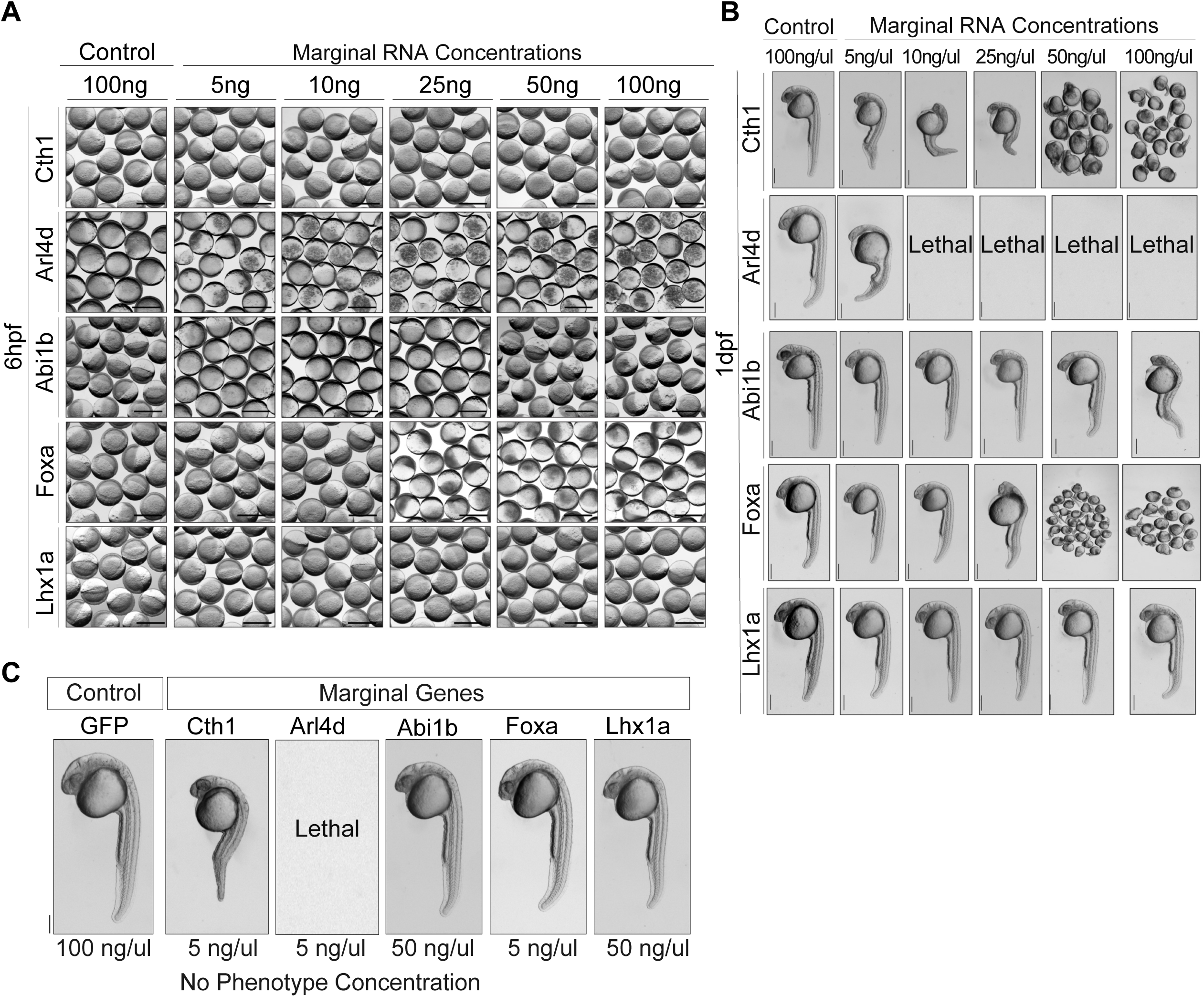
Embryos show different levels of tolerance to different ectopic concentration of marginal RNAs. A. Representative images of embryos at 6 hpf injected with open reading frame (ORF) RNA concentrations (5pg/embryo to 100pg/embryo) for respective marginal genes (*cth1, arl4d, abi1b, foxa* and *lhx1a*), as a control GFP mRNA (100pg/embryo) was used. Scale bar, 1000μm. B. Representative images of embryos at 26 hpf injected with open reading frame (ORF) RNA concentrations (5pg/embryo to 100pg/embryo) for respective marginal genes (*cth1, arl4d, abi1b, foxa* and *lhx1a*), as a control GFP mRNA (100pg/embryo) was used. Different marginal RNA shows different levels of respective ectopic RNA tolerance (*abi1b* (50pg/embryo), *foxa* (5pg/embryo), *lhx1a* (50pg/embryo)) (C), while in some cases minimal ectopic RNA concentration injected was also causing developmental deformities (*cth1* and *arl4d*, 5pg/embryo). Scale bar 0.5mm (single image panel). Images with multiple embryos are at different scale.

**Supplementary Figure 3.**
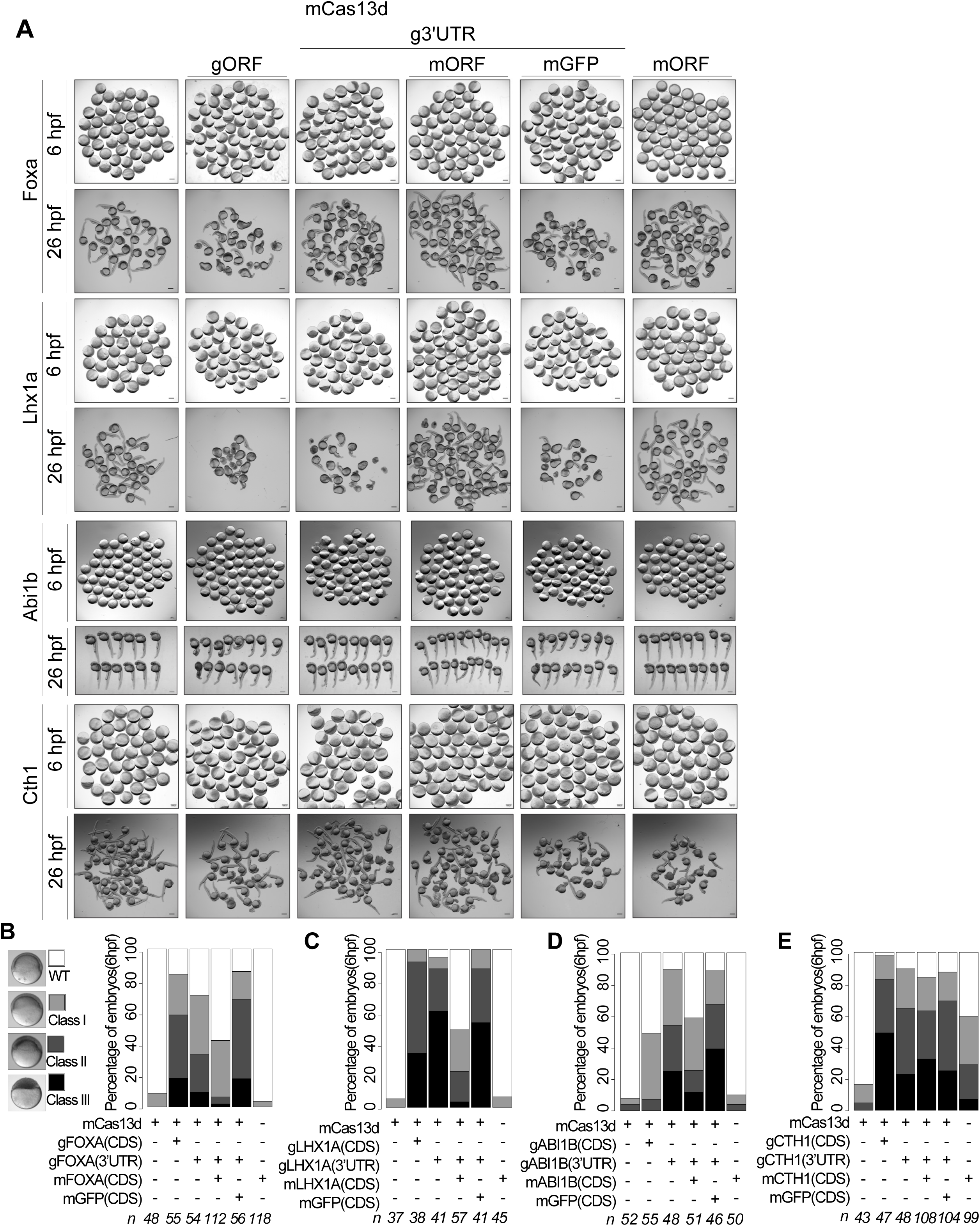
CRISPR-RfxCas13d mediated knockdowns are specific and can be rescued. A. Knockdown phenotype rescue experiments for marginal genes *foxa, lhx1a, abi1b* and *cth1* at 6 hpf and 1 dpf. Representative images of embryos injected with indicated molecules. Panel of mCas13d along with guideRNA target the endogenous 3’UTR (g3’UTR) and ectopic mRNA encoding for the marginal gene (mORF) is for phenotype knockdown rescue while other panels are different type of controls to respective rescue experiment for different marginal genes at 6 and 26 hpf. Scale bar, 1000 μm. B. C, D, E: Stack bar plots showing the percentage quantification of knockdown phenotype rescue for *foxa (B), lhx1a (C), abi1b (D)* and *cth1(E)* respectively at 6 hpf. Different intensities of gray color are positively associated with the severity of the phenotype (dark, most severe phenotype (class III) and white is wildtype). n represents the number of embryos observed (scale bar, 500 μm).

**Supplementary Figure 4.**
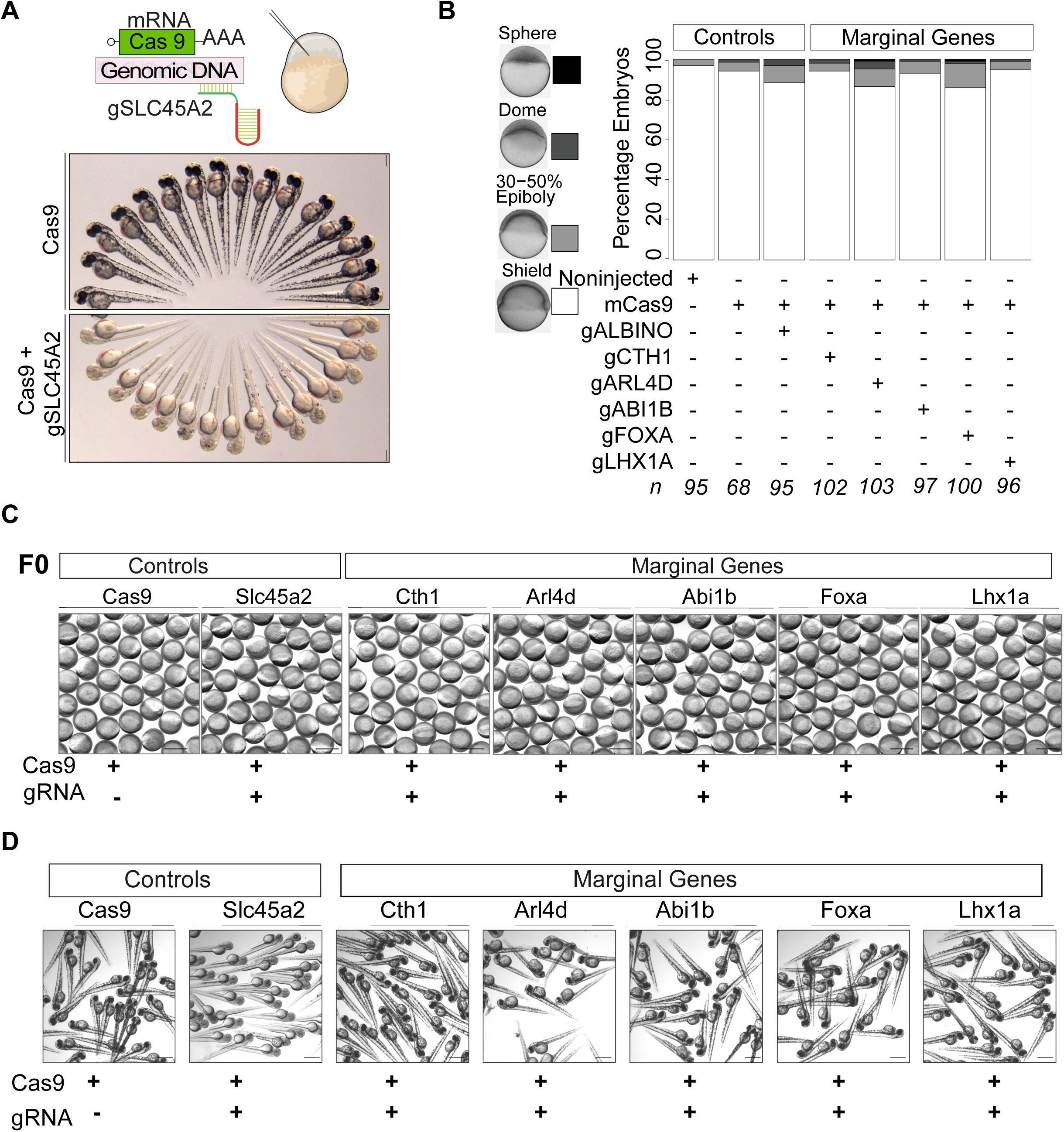
CRISPR-Cas9 mutation show minimal or no zygotic contribution on marginal gene phenotypes. A. CRISPR-Cas9 mutagenesis assay for *albino (Slc45a2)* gene. Upper panel shows Cas9 alone injected controls (100pg/embryo) larvae, lower panel show Cas9 (100pg/embryo) along with gRNA (25pg/embryo) against albino gene. Larvae at 2 dpf in albino targeted gene are either with no or minimal skin pigments while Cas9 controls are with high pigmentations. Scale bar, 2mm. B. Stack bar plot showing no significant Cas9 mediated mutation phenotypes for zygotic copies of marginal genes (*cth1, arl4d, abi1b, foxa* and *lhx1a*) as compared to different controls (noninjected, cas9 alone and albino) at 6 hpf. Different intensities of gray color are directly associated with severity of developmental phenotype (darkest shade for sphere, while white color is for shield stage). N represents the number of embryos observed. Scale bar, 500 μm. C. Embryos at 6 hpf and 2 dpf (D) from Cas9 mediated mutagenesis for different marginal genes (*cth1, arl4d, abi1b, foxa* and *lhx1a*) along with different controls (cas9 alone and albino) at 6 hpf. Embryos and larvae from Cas9 targeted marginal genes do not show severe phenotypes. Scale bar is 1000 μm for 6 hpf and 2mm for 2 dpf.

**Supplementary Figure 5.**
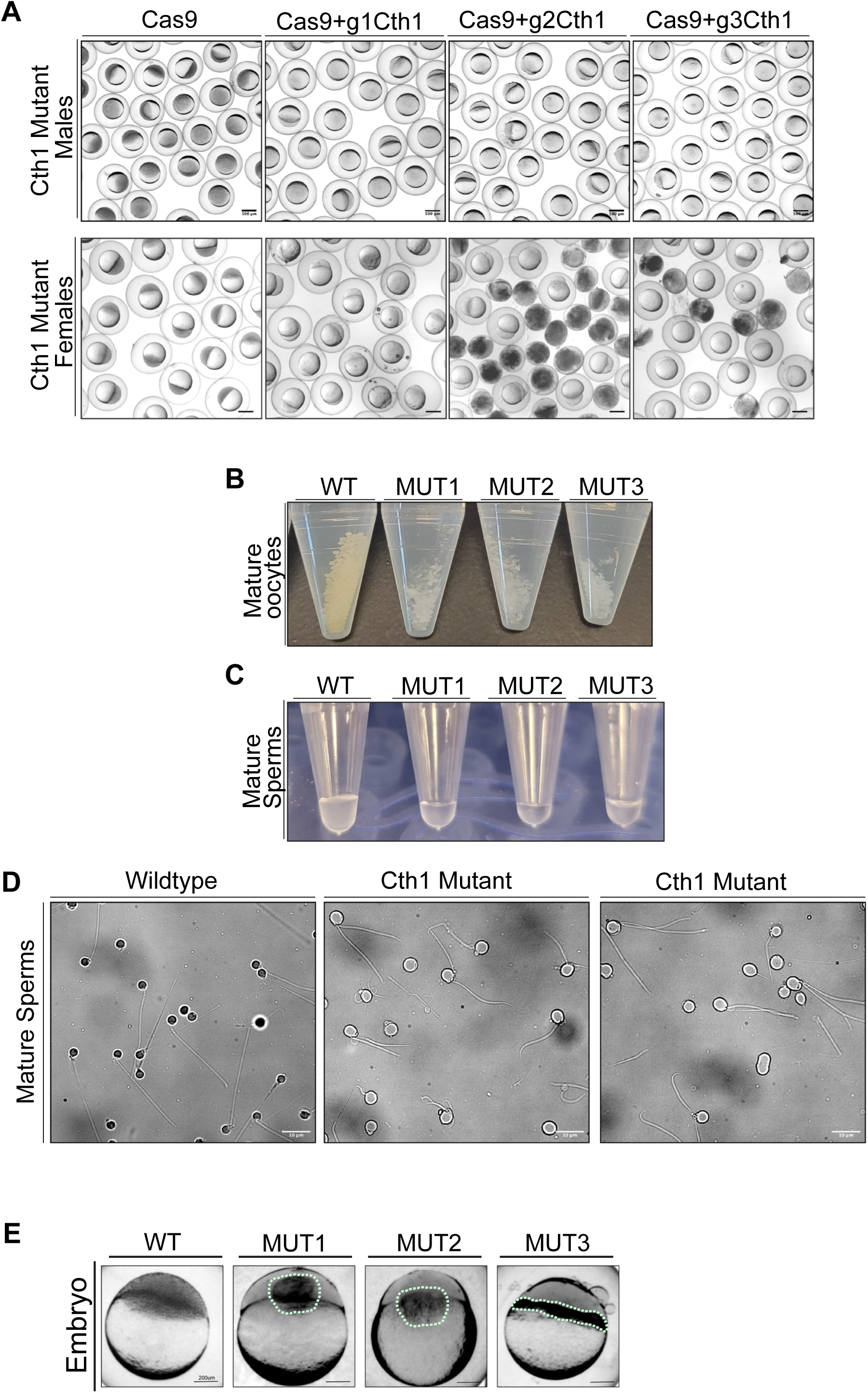
*Cth1* Cas9 mutants are infertile with abnormal mature gametes. A. Representative images of *cth1* mutant embryos resulted from the Cas9 mutagenesis with individual gRNA targeting *cth1* genomic locus. *Cth1* mutants when crossed with wildtype partner produce either unfertilized or degraded embryos. All the images are at 4hpf with scale bar 500 μm. B. Images of oocytes from three independent *cth1* female mutants are translucent white and degraded as compared to control oocytes which are intact, opaque yellow and round. Mature oocytes are in 15 ml conical tubes filled with 0.5X E2 (ZIRC) Embryo Media. C. Images of sperm squeezed from three independent *cth1* mutant and one wildtype adult zebrafish males. Tubes containing wildtype sperm are more turbid, and milky white while tubes with mutant sperm are less turbid and watery. These tubes are 0.2 ml PCR tubes filled with sperm storage buffer. D. Images of sperm from two independent *cth1* mutant males have abnormal morphology ranging from only head, multiple tails or fragment tails as compared to control intact head and one tail sperm squeezed from wildtype male of same age (14-month-old). Scale Bar 10 μm, Magnification 100X. E. Representative images of embryos at 4 hpf from *cth1* mutant males when bred with same age wildtype female. Embryos from the control group are at sphere stage, while embryos produced by crossing mutant males and wildtype females are unfertilized showing the unsuccessful process of fertilization. All the embryos are at 4hpf with scale bar 200 μm.

**Supplementary Figure 6.**
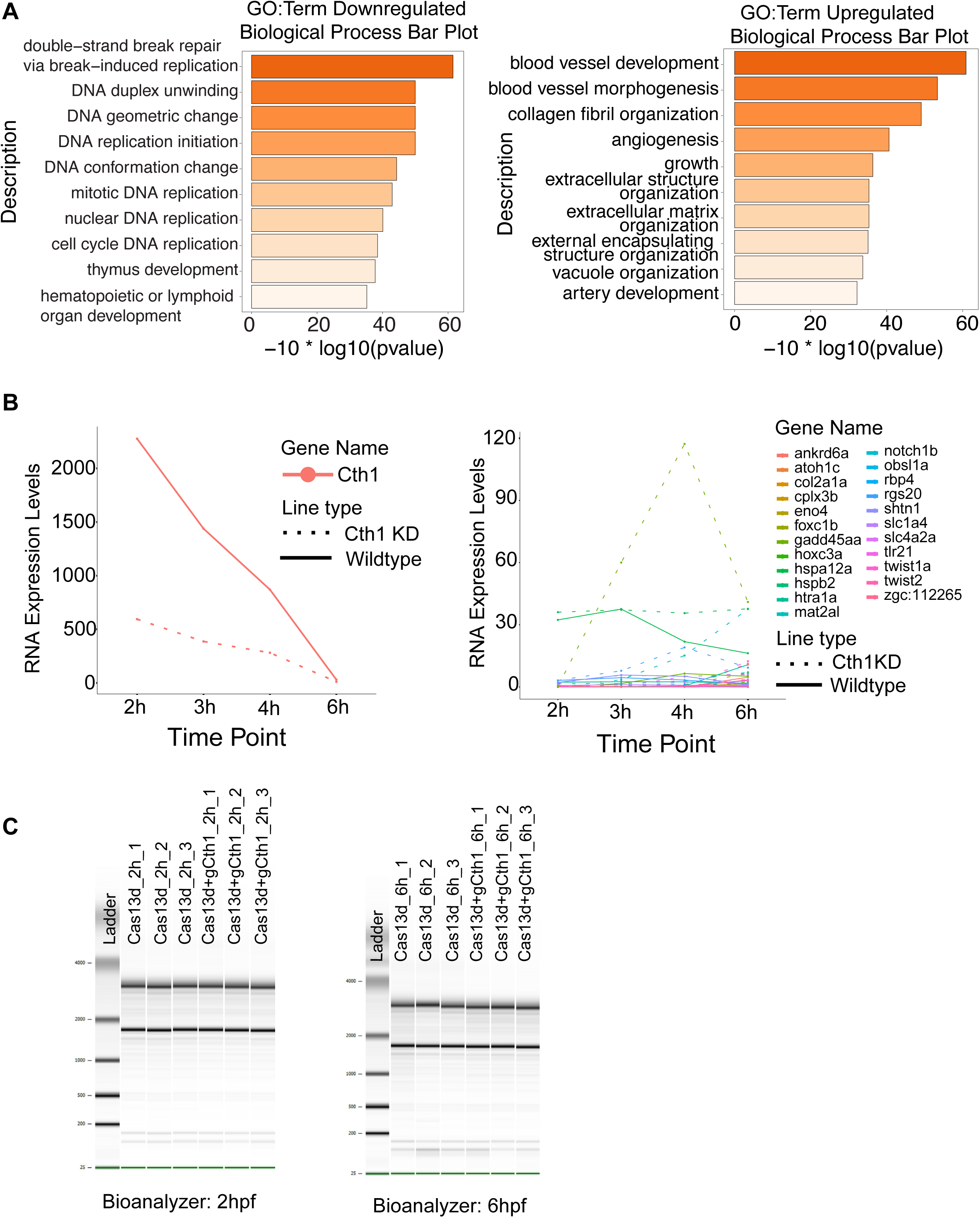
CRISPR-Cas13d uncover the functions of embryonic lethal *Cth1* gene during development. A. GO terms associated with Cas13d mediated *cth1* knockdown: First panel bar plot, GO terms for biological processes from differentially downregulated genes are mainly associated with DNA replication, repair and DNA conformational changes. Second panel bar plot, GO terms are differentially upregulated genes are mainly associated with blood, blood tissues, angiogenesis. B. Line plot showing the time course RNA-seq in wildtype and *cth1* knockdown conditions: RNA levels of *cth1* (first panel) and its potential targets (second panel) in wildtype conditions (solid line) and knockdown conditions (dashed line) at different time point (2 – 6 hpf). Upon *cth1* knockdown, its potential targets are getting upregulated (respective color dashed line). C. Bioanalyzer gels for 2 and 6 hpf RNA seq for cas13d alone (controls) and Cas13d + gcth1 (*cth1* knockdowns) showing no extra band between 28s and 18s rRNAs, which suggest no cas13d mediated collateral activity upon *cth1* knockdowns.

